# Advanced eTAM-seq enables high-fidelity, low-input *N*^6^-methyladenosine profiling in human cells and embryonic mouse tissues

**DOI:** 10.1101/2025.05.11.653357

**Authors:** Emily Mengshu He, Yuan Wu, Chang Ye, Tie-Bo Zeng, Chuan He, Weixin Tang

**Author notes:** These authors contributed equally (Emily Mengshu He, Yuan Wu, Chang Ye).

## Abstract

Functional dissection of N6-methyladenosine (m^6^A), the most abundant internal messenger RNA modification in mammals, demands quantitative, scalable detection technology. We previously reported eTAM-seq, which supports transcriptome-wide quantification of m^6^A by enzyme-assisted adenosine deamination. While effective, TadA8.20—the enzyme used in our first-generation technology—is sensitive not only to m^6^A but also to RNA structure, making accurate detection dependent on the inclusion of control transcriptomes. Here, we introduce eTAM-seq-v2, in which we replace TadA8.20 with TadA8r, a further evolved adenosine deaminase with superior catalytic efficiency. eTAM-seq-v2 supports control-free m^6^A calling with high fidelity. Because enzyme treatment preserves RNA integrity, eTAM-seq surveys >51% of A sites in all expressed genes with moderate sequencing depth (60 million uniquely mapped reads) and delivers robust performance with as little as 10 ng of total RNA (∼500 cells). With eTAM-seq-v2, we delineate the m^6^A landscape across six human cell lines and seven embryonic mouse tissues. While uncovering broadly conserved m^6^A patterns, we reveal that most neighboring m^6^A sites are independently deposited at the single-molecule level. Moving forward, we envision that eTAM-seq-v2 will enable researchers to survey m^6^A in diverse biological contexts and uncover new insights into its regulatory roles.

## Introduction

Chemical modifications have a profound impact on RNA function^1–4^. Among them, *N*^6^-methyladenosine (m^6^A) is the most abundant internal modification in mammalian mRNA. Its multifaceted roles in physiology and pathology^5, 6^ have been established by various profiling methods, many of which relied on m^6^A-specific antibodies^7, 8^. Recently, we reported evolved TadA-assisted *N*^6^-methyladenine sequencing (eTAM-seq)^9^, a method that locates and quantifies m^6^A via TadA8.20-assisted global A deamination. Around the same time, glyoxal and nitrite-mediated deamination of unmethylated adenosines (GLORI), a chemical deamination-based m^6^A detection method, was also reported^10, 11^. Since their introduction, these quantitative, base-resolution detection approaches have rapidly advanced our understanding of the regulatory networks shaped by m^6^A^12, 13^.

Our first-generation technology, eTAM-seq-v1, is sensitive not only to m^6^A but also to local RNA structure, and thus requires matched control transcriptomes for accurate detection. This requirement adds complexity to the workflow, limits detection sensitivity, and necessitates sophisticated biostatistical analysis. In this study, we introduce eTAM-seq-v2, an updated version of our method that features a further evolved deaminase^14^. With superior deamination efficiency, eTAM-seq-v2 eliminates the need for control transcriptomes and increases both detection sensitivity and fidelity. Additionally, we present a low-input eTAM-seq workflow that supports robust m^6^A calling with as little as 10 ng of total RNA.

Despite its well-documented, widespread regulatory roles, high-resolution, quantitative maps of m^6^A remain scarce^15, 16^, posing a barrier to functional interrogation. Another important goal of this study is to generate high-quality datasets for human cell lines and mouse tissues, providing a comparative, quantitative view of the m^6^A landscape. Together, these efforts not only expand the toolkit for m^6^A detection but also advance our molecular understanding of this essential epitranscriptomic mark.

## Results

### TadA8r is more active than TadA8.20 in deaminating RNA

In the original eTAM-seq workflow (eTAM-seq-v1), we chose TadA8.20—a hyperactive adenosine deaminase evolved for base editing^17^—to facilitate global A deamination for m^6^A detection. Since then, our lab has evolved TadA8r, which demonstrates increased potency and R<u>A</u> (R = A or G) compatibility on DNA substrates^14^. As most m^6^A appears in DRACH motifs (D = A, G, or U; H = A, C, or U), we hypothesized that the increased potency and broadened context compatibility of TadA8r might be particularly beneficial for m^6^A detection in eTAM-seq. We therefore set out to characterize the deamination capacity of TadA8r on RNA substrates and compare it with that of TadA8.20.

We incubated an 8-nucleotide (nt) RNA probe containing a single A with TadA8r or TadA8.20 at different concentrations for 1 h and analyzed the reactions using liquid chromatography-mass spectrometry (LC-MS; **Fig. 1a**). While 8.75 μM TadA8r fully deaminated the RNA substrate, complete deamination by TadA8.20 required a 4-fold higher enzyme dose (35 μM). Correspondingly, the half maximal effective concentration (EC_50_) for TadA8.20 was determined to be 7.6 μM, 3.3-fold higher than that of TadA8r (2.3 μM; **Fig. 1b**).

**Figure 1.**
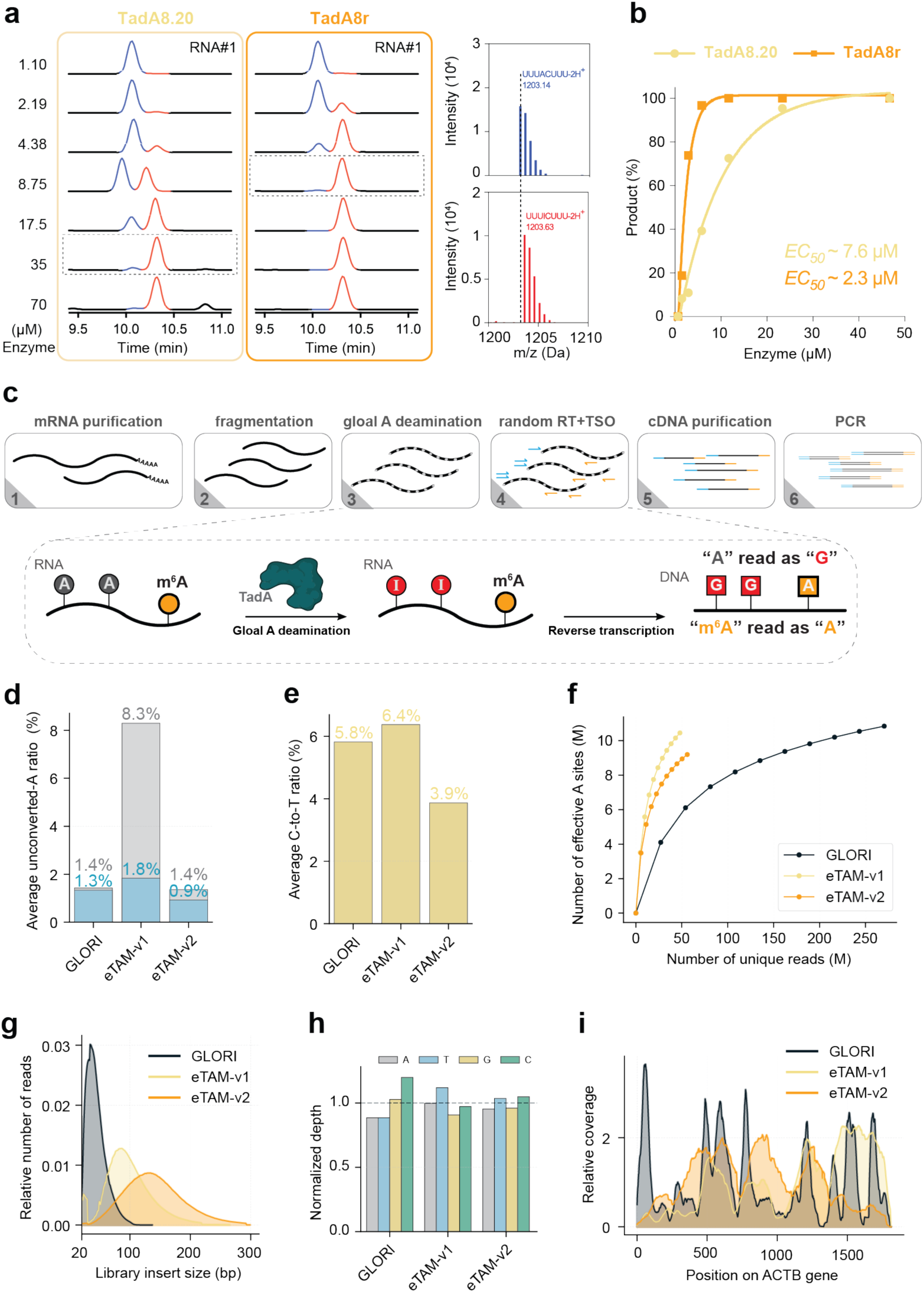
Performance of eTAM-seq-v2. **a**. TadA8.20- and TadA8r-mediated A deamination in an 8-nt RNA probe. RNA was incubated with different concentrations of the enzyme at 44 °C for 3 h and analyzed by LC-MS. Extracted ion chromatograms are shown, with representative mass spectra of the starting material and product displayed on the right. **b**. Half-maximal effective concentrations (EC_50_) for TadA8.20 and TadA8r. The conversion rates in **a** were quantified based on standard curves constructed using known concentrations of the starting material and product and fitted using a nonlinear regression model in GraphPad. **c**. Schematic of eTAM-seq-v2. RT: reverse transcription; TSO: template switch oligo; cDNA: complementary DNA. **d**. Average unconverted A rates detected in GLORI, eTAM-seq-v1, and eTAM-seq-v2 before (grey) and after (blue) removing reads with three or more consecutive unconverted As. **e**. Average C-to-T mutation rates in GLORI, eTAM-seq-v1, and eTAM-seq-v2. Mutation rates were calculated using uniquely mapped reads without filtering for A conversion. **f**. Number of effective A sites detected in GLORI, eTAM-seq-v1, and eTAM-seq-v2 under different sequencing depths. Effective A sites were defined as those with a sequencing depth >10. To avoid technical variations introduced during library construction, uniquely mapped reads were used to generate the plot. **g**. Distribution of insert lengths in GLORI, eTAM-seq-v1, and eTAM-seq-v2. **h**. Normalized base composition detected in GLORI, eTAM-seq-v1, and eTAM-seq-v2. **i**. Relative coverage across the *ACTB* transcript in GLORI, eTAM-seq-v1, and eTAM-seq-v2. Coverage at individual positions was normalized to the coverage of the entire transcript. Plots were generated using published GLORI data from HEK293T mRNA (SRR21356251) and eTAM-seq data from HeLa mRNA.

Consistent with our previous studies^14^, TadA8r outperformed TadA8.20 on DNA probes (**Extended Data Fig. 1a-b**). When heated, both proteins remained folded up to 56-59 °C (**Extended Data Fig. 1c**), in line with activity assays indicating that TadA8r was only fully deactivated above 60 °C (**Extended Data Fig. 1d**). As RNA secondary structures can resolve at high temperatures, the heat-resistant property of TadA8r may help reduce the background caused by unconverted As in structured regions, which was a major challenge during the development of eTAM-seq-v1. Incubation of TadA8r with total RNA leaves the RNA intact (**Extended Data Fig. 1e**), confirming that enzymatic conversion preserves RNA integrity. Collectively, these findings demonstrate that TadA8r outperforms TadA8.20 in global A deamination, remains active at elevated temperatures, and maintains RNA integrity.

### Benchmarking eTAM-seq-v2

We incorporated TadA8r into an eTAM-seq workflow to develop eTAM-seq-v2 (**Fig. 1c** and **Supplementary Table 1**). In addition to replacing the enzyme, we switched to a commercial RNA library construction kit to facilitate facile adaptation of our technology (see **Methods**). When applied to 50 ng of HeLa poly(A)^+^ RNA (hereafter referred to as HeLa mRNA), eTAM-seq-v2 delivered a global deamination rate of 98.6%, measured as the fraction of G counts relative to the combined A and G counts mapped to genomic A sites, which was higher than the 91.7% achieved by eTAM-seq-v1 and comparable to the 98.6% obtained by GLORI^10^ (data from HEK293T samples prepared by a ligation-based method; **Fig. 1d**). We further filtered out reads with three or more consecutive unconverted As to mask poorly processed RNA fragments, which increased the global deamination rates of GLORI, eTAM-seq-v1, and eTAM-seq-v2 to 98.7%, 98.2%, and 99.1%, respectively (**Fig. 1d** and **Extended Data Fig. 2a**).

C deamination is a common side reaction that occurs during TadA- and glyoxal/nitrite-mediated A deamination^10, 11, 17^. While achieving a higher global A deamination rate, eTAM-seq-v2 showed markedly lower C-to-U conversion (3.9%) compared to eTAM-seq-v1 (6.4%; **Fig. 1e** and **Extended Data Fig. 2b)**. Interestingly, C-to-U conversion in GLORI demonstrated strong batch-to-batch variation (5.8-15.8%; **Fig. 1e** and **Extended Data Fig. 2b)**, suggesting that side reactions may be more challenging to control during chemical deamination. As excessive C-to-T mutations can compromise mapping, eTAM-seq-v2 is expected to generate a higher proportion of mappable reads.

We next examined the number of effective A sites—defined as sites with sequencing depth greater than 10— which largely determines the number of detectable m^6^A sites, relative to the number of uniquely mapped reads across the three sequencing methods (**Fig. 1f**). With 50 million uniquely mapped reads, eTAM-seq-v1 and eTAM-seq-v2 covered 8–10 million A sites, while GLORI required 200 million reads to achieve comparable A-site detection. This discrepancy is expected, as eTAM-seq-v1 and eTAM-seq-v2 generated average insert lengths of 86 and 134 nt, respectively, much longer than GLORI inserts (34 nt; **Fig. 1g** and **Extended Data Fig. 2c**).

We also noticed a bias in base composition, particularly an overrepresentation of C bases, in GLORI reads (**Fig. 1h** and **Extended Data Fig. 2d**), which prompted us to assess the uniformity of coverage across transcripts. Taking housekeeping genes—*ACTB*, *HPRT1*, *RPLP0*, *RPL13A*, and *GAPDH*—as examples, relative sequencing depth and coverage remained stable for eTAM-seq-v1 and eTAM-seq-v2 (**Fig. 1i** and **Extended Data Fig. 2e-h**). eTAM-seq-v1 showed slightly better coverage of the 3’ untranslated region (UTR), likely due to its ligation-based library construction. GLORI, in contrast, experienced greater variation in sequencing depth along transcripts (**Fig. 1i** and **Extended Data Fig. 2e-h**). We suspect that this variability arises from incomplete G deprotection, as glyoxal-protected G, an intermediate of GLORI, interferes with reverse transcription^18^. Collectively, eTAM-seq-v2 achieves a high global deamination rate, is less prone to C deamination, resists premature termination during reverse transcription, and supports even transcriptome coverage. These favorable parameters set the stage for faithful and sensitive m^6^A detection by eTAM-seq-v2.

### m^6^A detection by eTAM-seq-v2

We next attempted to call m^6^A from effective A sites (i.e., sequencing depth >10). We previously minimized false positive signals caused by structured A using control transcriptomes^19^—generated either by a demethylase, fat mass- and obesity-associated protein (FTO), or *in vitro* transcription (IVT). With improved global A deamination, we hypothesized that control-free m^6^A detection would be feasible for eTAM-seq-v2. We first plotted persistent A rates for 16 individual N<u>A</u>N motifs (**Fig. 2a-b**). While G<u>A</u>C and A<u>A</u>C, motifs hosting the majority of m^6^A sites^20^, contributed most persistent A signals, C<u>A</u>G and G<u>A</u>G also showed average signals above 2%, which may introduce non-specific background.

**Figure 2.**
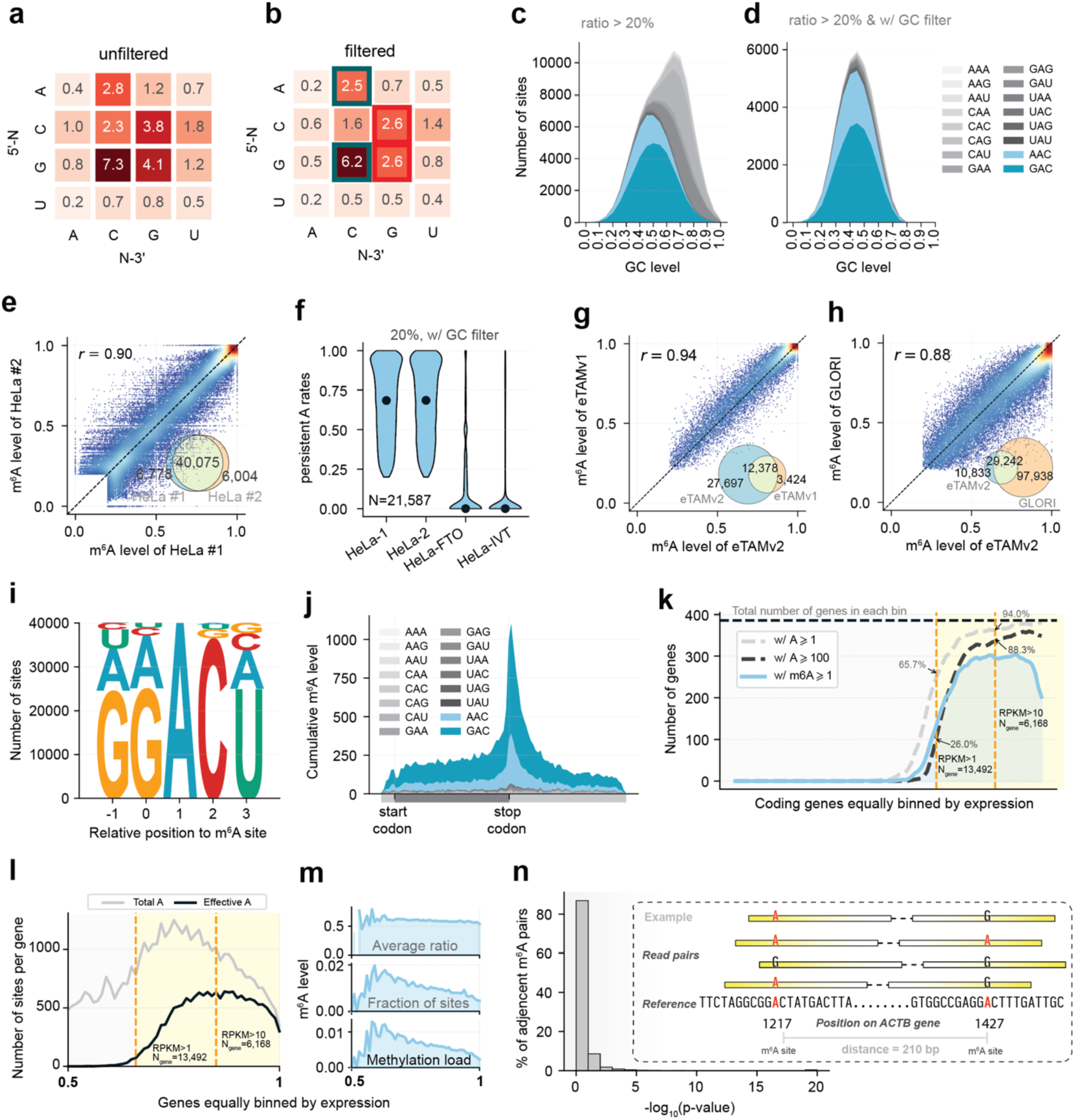
m^6^A profiling in HeLa mRNA by eTAM-seq-v2. **a**. Persistent A rates across 16 N<u>A</u>N contexts in unfiltered reads. **b**. Persistent A rates across 16 N<u>A</u>N contexts after removing reads with three or more consecutive unconverted As. **c**. Distribution of persistent A sites across different GC content without a GC filter. GC content was calculated using 10-nt upstream and 10-nt downstream sequences flanking the target A. A sites are colored based on their N<u>A</u>N sequence contexts, and those with ≥20% persistent A signals are plotted. **d**. Distribution of persistent A sites across different GC content after applying a GC filter. More information on the GC filter is provided in **Supplementary Note 1**. **e**. Methylation level correlation of m^6^A sites detected in two biological replicates. A sites with sequencing depth >10 in both replicates were considered. A Venn diagram showing the overlap of detected sites is inserted. **f**. Violin plot for methylation levels of overlapping m^6^A sites detected in two eTAM-seq-v2 replicates, FTO-treated HeLa mRNA, and an IVT HeLa transcriptome. All m^6^A sites with sequencing depth >1 in both FTO and IVT samples are plotted. Dots mark the median level of each sample. **g**. Methylation level correlation of m^6^A sites detected by eTAM-seq-v1 and eTAM-seq-v2. A Venn diagram showing the overlap of detected sites is inserted. m^6^A sites co-discovered by eTAM-seq-v1 replicates 1 and 2 were used. **h**. Methylation level correlation of m^6^A sites detected by GLORI and eTAM-seq-v2. A Venn diagram showing the overlap of detected sites is inserted. GLORI data were generated using HEK293T mRNA; m^6^A sites co-discovered by GLORI replicates 1 and 2 were used. **i**. Logo plot of the consensus sequences hosting m^6^A. **j**. Metagene plot of transcriptome-wide distribution of m^6^A. **k**. Number of genes with m^6^A and 1 or 100 effective A sites binned by expression percentile. All 38,549 ENCODE-annotated genes (protein_coding and lncRNA) are binned by expression. RPKM: reads per kilobase per million mapped reads. **l**. Average effective and total A sites per gene across different expression levels. **m**. Average methylation stoichiometry per m^6^A, m^6^A frequency per effective A, and methylation load per gene across different expression levels. **n**. Co-occurrence of adjacent m^6^A sites. A diagram of an m^6^A pair and spanning reads is inserted. m^6^A sites co-detected in two eTAM-seq-v2 replicates are plotted in panels **g**-**j**. Replicate 1 was analyzed for **k**-**n**.

We considered three potential causes of conversion resistance: 1) enzyme context preferences, 2) local secondary structure, and 3) differential amplification of A- and I-containing strands during reverse transcription and subsequent PCR. Although TadA8r is globally more active than TadA8.20 across all N<u>A</u>N motifs, minor deficiencies were observed at C<u>A</u>G and G<u>A</u>G motifs. Interestingly, this context preference differs from that observed in DNA—prior studies have shown that TadA8r is least efficient at processing A<u>A</u>A in DNA substrates^14^—suggesting that the enzyme may engage RNA and DNA through distinct binding modes. Both the second and third factors may correlate with local GC content. Indeed, we observed an exponential increase in conversion resistance with higher GC content within 20 nt of the target A for all sequence contexts (**Extended Data Fig. 3a**). It is worth noting that most A sites have balanced GC content (∼50%) in this 20-nt window (**Extended Data Fig. 3b**), meaning that high local GC content impacts only a small fraction of all A sites. Based on this trend, we modeled conversion resistance as a function of local GC content and subtracted this background to facilitate m^6^A calling (**Fig. 2c-d** and **Supplementary Note 1**).

We detected 61,100 and 46,868 m^6^A sites with methylation levels equal to or more than 20% in two biological replicates (60/38 million uniquely mapped reads; **Supplementary Table 2**), of which 40,075 were shared (85.5% and 87.0% considering sites effectively surveyed in both replicates; **Fig. 2e**). Methylation levels were highly consistent between replicates (Pearson’s *r* = 0.90), with a mostly even distribution and a slight enrichment toward 100% (**Fig. 2e**). The correlation coefficient increased to 0.98 when depth cutoff was raised to 100 (**Extended Data Fig. 3c-e**), indicating that variations in methylation quantification primarily stem from insufficient sequencing depth of lowly abundant transcripts. To evaluate the fidelity of eTAM-seq-v2, we prepared an FTO-treated HeLa transcriptome and an *in vitro* transcribed, modification-free HeLa transcriptome^19^. The median methylation levels of our detected m^6^A sites dropped from 68-69% in two replicates to 0% in both FTO-treated and IVT samples (**Fig. 2f**), confirming that these were genuine m^6^A sites. Similar trends were observed for sites with 10-20% methylation (**Extended Data Fig. 3f**). Further, eTAM-seq-v2-reporpted methylation levels were consistent with those of eTAM-seq-v1 (Pearson’s *r* = 0.94; **Fig. 2g**) and GLORI results (Pearson’s *r* = 0.88, HEK293T mRNA; 127,180 m^6^A sites with ≥20% methylation detected using 390 M uniquely mapped reads; **Fig. 2h** and **Extended Data Fig. 3g-h**). As expected, m^6^A primarily appeared in DRACH motifs (**Fig. 2i**) and was enriched in 3’ UTRs near the stop codon (**Fig. 2j**). Together, these results indicated that eTAM-seq-v2 enables reliable m^6^A detection and quantification.

### Transcriptome coverage and co-occurrence of m^6^A

To inspect the transcriptome coverage, we analyzed gene expression using eTAM-seq reads, which have previously been shown to correlate well with RNA-seq data^9^. Among the 38,549 protein-coding and long non-coding RNA (lncRNA) genes annotated by the Encyclopedia of DNA Elements (ENCODE)^21, 22^, 13,492 and 6,168 were expressed at levels above 1 and 10 reads per kilobase per million mapped reads (RPKM), respectively—corresponding to low and moderate expression. The remaining detected genes were deemed very lowly expressed. We binned all genes by expression percentile and examined the number of effective A sites in each bin (**Fig. 2k**). Among genes with RPKM >1, 65.7% had at least one effective A site, a proportion that increased to 94.0% among genes with RPKM >10. In fact, more than 88.3% of genes with RPKM >10 had at least 100 effective A sites. Although having more effectively covered A sites is expected to promote m^6^A detection, the proportion of m^6^A-containing genes in each bin appeared saturated at moderate expression levels and unexpectedly declined at higher expression levels.

To systematically examine the declining frequency of m^6^A-containing genes, we plotted the numbers of total and effective A sites per gene across exonic regions within each expression bin (**Fig. 2l**). The number of effective A sites per gene increased and peaked at ∼600 as expression rose from RPKM 0 to 10. This is expected since sequencing coverage directly dictates effective A site counts. The number of effective and total A sites per gene gradually converged among highly expressed genes (RPKM >10), indicating that detection was approaching saturation. We note that the number of total A sites declined significantly in highly expressed genes, consistent with their shorter transcript lengths^23, 24^. Taken together, these findings suggest that, at our sequencing depth (60 million uniquely mapped reads), we captured >51% of A sites in all expressed genes (n = 13,492; RPKM >1), and that detection approached saturation for moderately and highly expressed genes (n = 6,168; RPKM >10). Although deeper sequencing is expected to further boost m^6^A detection, additional gains will likely come predominantly from genes with lower expression.

We next inspected m^6^A detection across expression bins (**Fig. 2m**). While the average methylation level remained steady (50-60%), the frequency of m^6^A sites per effective A site consistently decreased as expression levels increased. Consequently, there was a clear downward trend in methylation load per gene— defined as 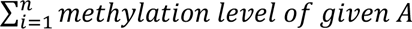 / total number of effective A sites—as a function of expression levels. These findings suggest that highly expressed genes tend to be less methylated, a subtle effect recently noted in immunoprecipitation-based and single-nucleotide resolution m^6^A analyses^25, 26^.

We and others have previously demonstrated that m^6^A sites can cluster within the same gene^9, 10, 27–29^. Recently, several direct RNA sequencing studies have suggested a tendency for m^6^A to co-occur in the same transcript^30, 31^. While able to cover the entire transcript, direct RNA sequencing is currently less accurate in m^6^A detection. In contrast, conversion-based detection methods, such as eTAM-seq and GLORI, offer more reliable m^6^A calling but do not preserve long-range crosstalk signals. Nevertheless, with an average insert length of 134 nt, eTAM-seq-v2 retains co-occurrence information of nearby m^6^A sites, i.e., persistent A signals in the same read. To quantify co-occurrence, we screened all 308,567 m^6^A pairs derived from 55,830 sites identified in exonic regions. Analysis of 34,694 m^6^A pairs, located up to 300 nt apart and covered by at least 10 reads, revealed that 33,123 (95%) exhibited an independent profile (*P* ≥ 0.01; **Methods**; **Fig. 2n** and **Extended Data Fig. 4a-b**). These findings are cross validated using GLORI data (99,226 m^6^A sites with a modification ratio ≥20%), although fewer m^6^A pairs were detected due to its shorter read length (**Extended Data Fig. 4c-d**). Collectively, these results suggest that the deposition of most neighboring m^6^A sites is independent.

### m^6^A landscape in six human cell lines

We subjected six human cell lines derived from various tissues—HEK293T (embryonic kidney), HeLa (cervix), HepG2 (liver), Jurkat (T lymphocytes in blood), K562 (lymphoblasts in bone marrow), and U87 (brain)—to eTAM-seq-v2 in two biological replicates (**Fig. 3a**). Across these 12 samples, we detected 33,341-76,402 m^6^A sites, 83-86% of which were in RAC motifs, with 3’ UTR enrichment observed consistently (**Fig. 3b** and **Supplementary Table 2**). Note that the number of m^6^A sites detected in each sample is influenced by sequencing depth—i.e., transcriptome coverage—rather than directly reflecting methylation density in a given transcriptome. In FTO-treated samples, only a few hundred sites were detected, whereas IVT controls yielded thousands, potentially due to conversion-resistant double-stranded RNA generated during *in vitro* transcription^32^. Nevertheless, putative sites detected in control samples showed neither motif nor 3’ UTR enrichment (**Extended Data Fig. 5a**). Among detected m^6^A sites, methylation stoichiometry was higher at G<u>A</u>C than at other N<u>A</u>N motifs (**Extended Data Fig. 5b**), consistent with the G<u>A</u>C preference of the METTL3-METTL14 writer complex previously observed both *in vivo*^33^ and *in vitro*^28, 34^. m^6^A levels detected across different transcript regions—5’ UTR, coding DNA sequence (CDS), and 3’ UTR—remained stable (**Extended Data Fig. 5c**). In addition to the consistent methylation levels observed within replicates (Pearson’s *r* = 0.89-0.91; **Extended Data Fig. 5d**), pairwise comparison across different samples also revealed high concordance in detected m^6^A sites and their methylation levels (Pearson’s *r* = 0.86-0.92, **Extended Data Fig. 5e**).

**Figure 3.**
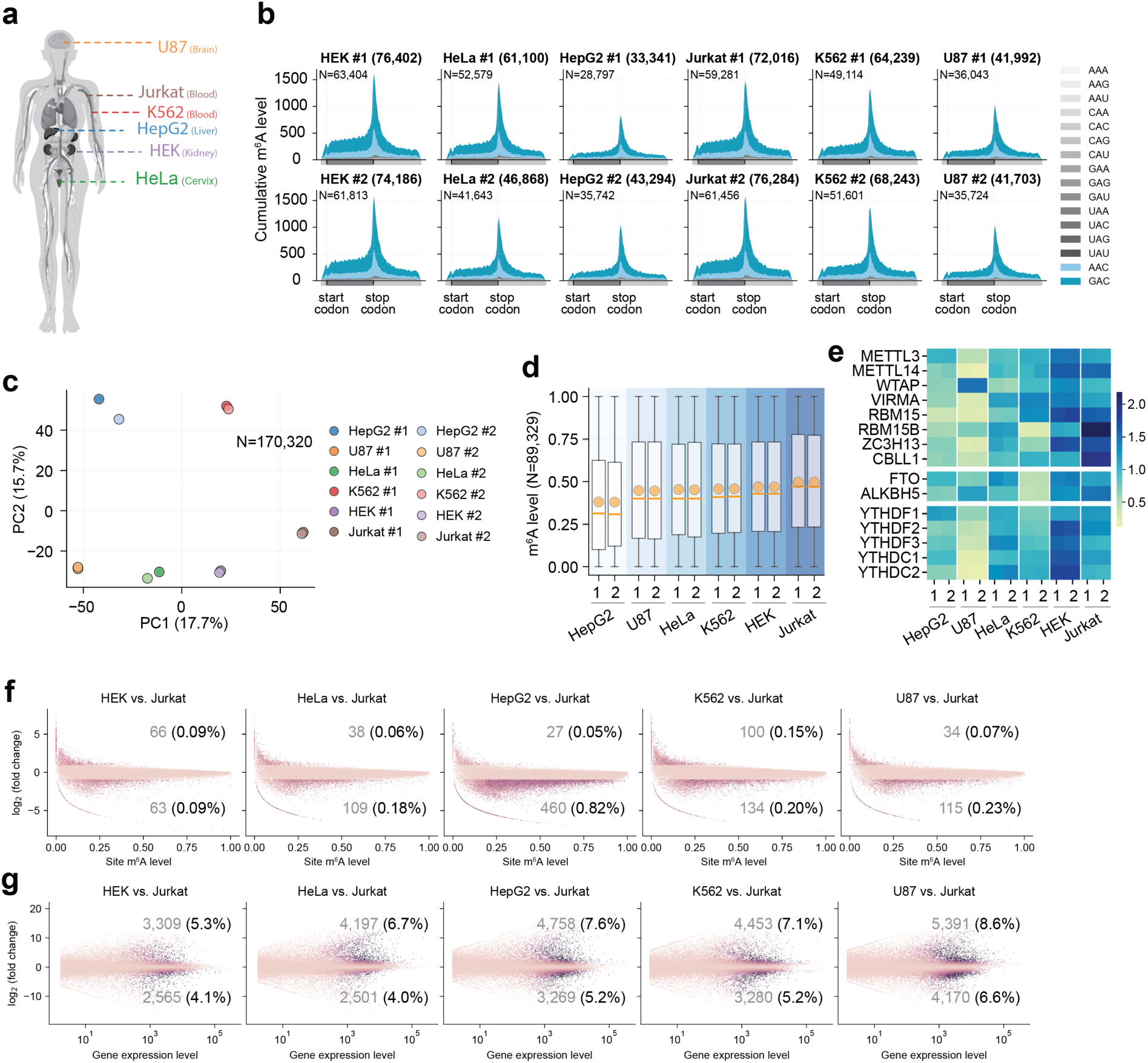
m^6^A landscape in six human cell lines. **a**. Diagram illustrating the origins of the six human cell lines analyzed using eTAM-seq-v2. Two biological replicates were conducted for each cell line. **b**. Metagene plot showing the distribution of detected m^6^A sites across mRNA transcript regions. The total number of detected m^6^A sites is indicated in the title, while the number of m^6^A sites (*N*) used for metagene analysis is noted within the panel. **c**. Principal component analysis (PCA) of m^6^A sites and their methylation levels across 12 samples. Sites identified as m^6^A in at least one sample were included in the analysis: n = 170,320. In this analysis, we allowed methylation levels to be calculated with <10 reads. Null value (A site without coverage in a given sample) was filled with 0. **d**. Box plot of global methylation levels detected in 12 samples. Sites identified as m^6^A in at least one sample and covered in all 12 samples (sequencing depth >0) were used to calculate global methylation levels: n = 89,329. Here we again allowed methylation levels to be calculated with <10 reads. **e**. Heatmap displaying expression levels of m^6^A writer, eraser, and reader proteins in six cell lines. **f**. Differential methylation between Jurkat and the other five cell lines. Differentially methylated sites are defined as those showing at least a two-fold difference in methylation levels between two cell lines and passing paired t-tests (*P* < 0.01). Sites identified as m^6^A in at least one sample and with an aggregated sequencing depth >20 in both cell types were included in the analysis. **g**. Differential expression analysis between Jurkat and the other five cell lines. Differentially expressed genes are defined as those with at least a two-fold difference in expression levels between two cell lines and passing paired Wald Tests in DESeq2 (adjusted *P* < 0.01). Methylation and expression levels in **f** and **g** were averaged from two biological replicates.

Despite the generally stable methylation patterns, biological replicates clustered together in principal component analysis (based on sites methylated in at least one sample; **Fig. 3c**), suggesting that m^6^A alone is sufficient to differentiate cell types. Among the six cell lines, Jurkat cells showed the highest global methylation level—calculated using sites methylated in at least one sample and covered in all samples (sequencing depth >0)—whereas HepG2 cells had the lowest. Specifically, the methylation level hierarchy was clearly defined as HepG2<U87<HeLa<K562<HEK293T<Jurkat (**Fig. 3d**). We correlated global methylation levels with the expression of core m^6^A writers *METTL3* and *METTL14*, as well as demethylases *ALKBH5* and *FTO*. Positive correlations were observed with *METTL3* and *METTL14* (Pearson’s *r* = 0.23 and 0.57; **Extended Data Fig. 5f**), suggesting that these writers regulate global m^6^A deposition. Surprisingly, expression of *FTO* and *ALKBH5* also showed weak but positive correlations with global methylation levels (Pearson’s *r* = 0.19 and 0.58; **Extended Data Fig. 5f**), indicating that demethylases play a minor role in shaping the m^6^A landscape and their impacts cannot be fully explained by their dedicated enzymatic activity. To further track down drivers of m^6^A homeostasis, we expanded our analysis and included additional components of the writer complex— *WTAP*, *VIRMA*, *RBM15*, *RBM15B*, *ZC3H13*, and *CBLL1*—as well as the reader family—*YTHDF1-3* and *YTHDC1-2*. Notably, the collective expression of writer components demonstrated a strong positive correlation with global methylation levels (**Fig. 3e**), suggesting that the whole writer complex, especially the regulatory subunits, dictates the m^6^A landscape. A positive correlation was also observed between global methylation levels and expression of reader proteins, albeit to a lesser extent (**Fig. 3e**).

We next investigated differential methylation among the six cell types. To maximize detection of cell type-specific methylation, we compared Jurkat, the cell line with the highest global methylation level, against the other five cell lines. We defined differentially methylated sites as those showing at least a two-fold difference in methylation levels between two cell lines. Additionally, paired t-tests were conducted on methylation levels measured from biological replicates of the two cell lines to minimize false positives (*P* < 0.01; **Methods**). Using these criteria, we detected 460, 115, 109, 134, and 63 differentially methylated sites with higher methylation levels in Jurkat compared to HepG2, U87, HeLa, K562, and HEK293T, respectively (**Fig. 3f**). Meanwhile, we identified 27, 34, 38, 100, and 66 sites with lower methylation levels in Jurkat relative to HepG2, U87, HeLa, K562, and HEK293T, respectively. These numbers align broadly with the global methylation levels detected in individual cell lines (**Fig. 3d**), suggesting that these differential methylation events are, at least in part, driven by proficiencies in global m^6^A deposition. As a control, we examined expression levels and confirmed that 4.0– 6.6% and 5.3–8.6% of transcripts were expressed at significantly higher and lower levels in Jurkat than in other cell types, respectively (**Fig. 3g**). Collectively, while differential methylation events are relatively infrequent, m^6^A alone appears sufficient to differentiate cell types.

### m^6^A landscape in mouse embryos

In addition to immortalized human cell lines, we explored the m^6^A landscape in mouse embryos. We harvested seven organs—brain, tongue, heart, lung, liver, kidney, and placenta—from two E18.5 mouse embryos (**Fig. 4a**). RNA was extracted separately from each organ and subjected to eTAM-seq-v2. Some organs, such as the kidney, were small and yielded lower-quality mRNA, resulting in reduced library quality and fewer detected m^6^A sites (**Supplementary Table 3**). Across these 14 samples (two biological replicates per organ), we detected 24,386-87,727 m^6^A sites, again enriched in RAC motifs and 3’ UTRs (**Fig. 4b** and **Extended data Fig. 6a-c**), with consistent methylation levels between replicates (Pearson’s *r* = 0.76-0.87; **Extended data Fig. 6d**).

**Figure 4.**
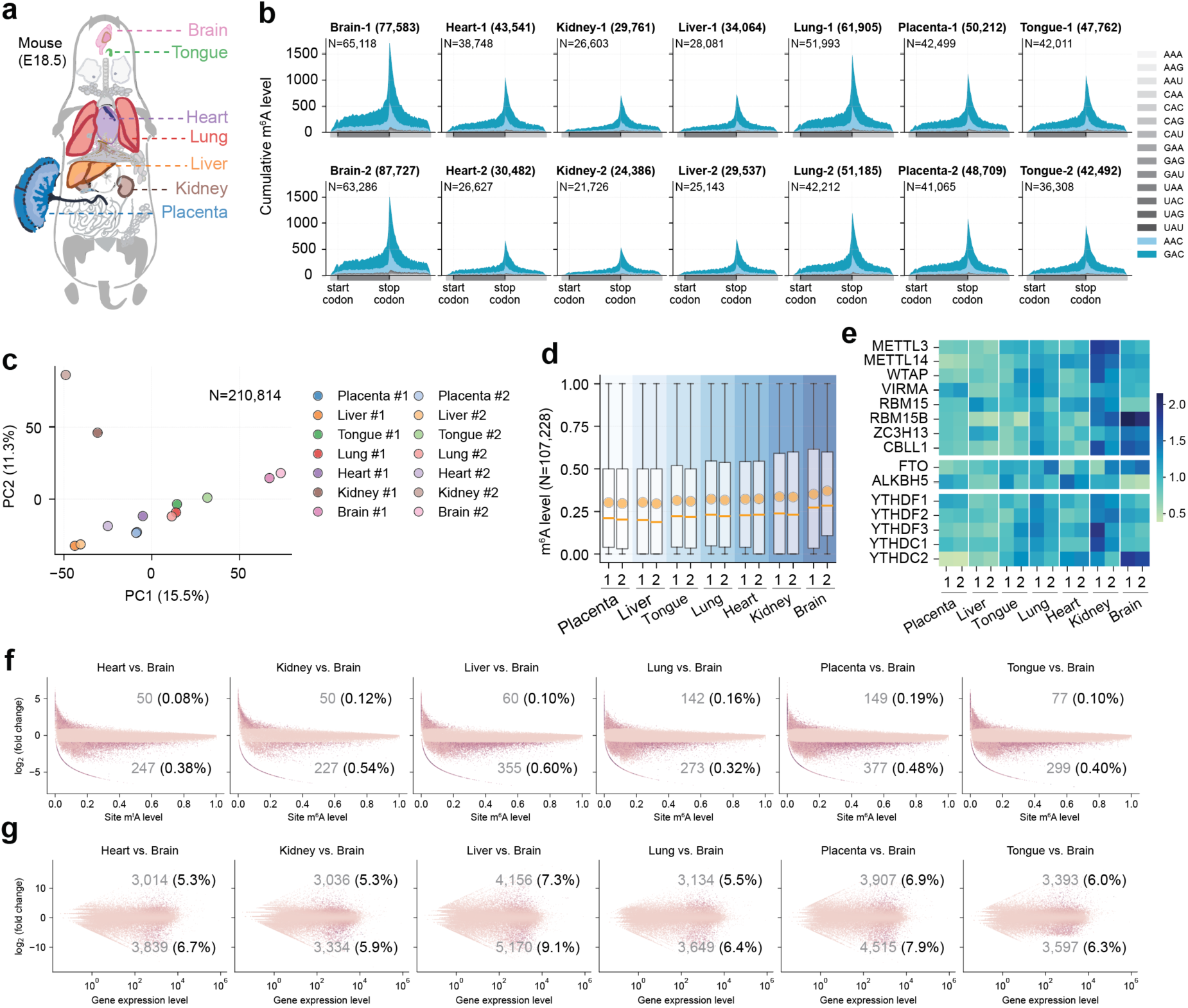
m^6^A landscape in mouse embryos. **a**. E18.5 mouse organs analyzed by eTAM-seq-v2. Two biological replicates were conducted for each organ. **b**. Metagene plot showing the distribution of detected m^6^A sites across mRNA transcript regions. The total number of detected m^6^A sites is indicated in the title, while the number of m^6^A sites (*N*) used for metagene analysis is noted within the panel. **c**. Principal component analysis (PCA) of m^6^A sites and their methylation levels across 14 samples. Sites identified as m^6^A in at least one sample were included in the analysis: n = 210,814. In this case, we allowed methylation levels to be calculated with <10 reads. Null value (A site without coverage in a given sample) was filled with 0. **d**. Box plot of global methylation levels detected in 14 samples. Sites identified as m^6^A in at least one sample and covered in all 14 samples (sequencing depth >0) were used to calculate global methylation levels: n = 107,228. Here we again allowed methylation levels to be calculated with <10 reads. **e**. Heatmap displaying expression levels of m^6^A writer, eraser, and reader proteins in seven mouse organs. **f**. Differentially methylated sites in mouse brain vs. heart, kidney, liver, lung, tongue, and placenta. Differentially methylated sites are defined as those showing at least a two-fold difference in methylation levels between two organs and passing paired t-tests (*P* < 0.01). Sites identified as m^6^A in at least one sample and with an aggregated sequencing depth >20 in both organs were included in the analysis. **g**. Differential expression between the brain and the other six organs. Differentially expressed genes are defined as those with at least a two-fold difference in expression levels between two organs and passing Wald Tests in DESeq2 (adjusted *P* < 0.01). Methylation and expression levels in **f** and **g** were averaged from two biological replicates.

As in human samples, pairwise comparison across mouse tissues also revealed largely consistent m^6^A sites and methylation levels (Pearson’s *r* = 0.87-0.92, **Extended Data Fig. 6e**). Principal component analysis again confirmed that m^6^A alone is sufficient to distinguish among tissues (**Fig. 4c**). Consistent with prior reports^35^, the brain had the highest methylation level among the seven organs (considering A sites methylated in at least one sample; **Fig. 4d**). Specifically, the organs ranked in increasing global methylation levels as follows: placenta<liver<tongue<lung<heart<kidney<brain. These global methylation levels correlated strongly with the collective expression of writer components, especially the regulatory subunits *RBM15B*, *ZC3H13*, and *CBLL1*, and to a lesser extent with the expression of reader proteins (**Fig. 4e** and **Extended Data Fig. 6f**).

We further investigated differential methylation events. We compared the brain—the organ with the highest global methylation level—against the other six organs. Using the same 2-fold difference and significance threshold, we identified 377, 355, 299, 273, 247, and 227 m^6^A sites with higher methylation levels in the brain compared to the placenta, liver, tongue, lung, heart, and kidney, respectively (**Fig. 4f**). Meanwhile, 149, 60, 77, 142, 50, and 50 sites were less methylated in the brain than in these organs. In contrast to the relatively infrequent differential methylation events, gene expression profiles varied considerably in these seven organs—5.9–9.1% and 5.3–7.3% of genes were expressed at significantly higher and lower levels in the brain than in other organs, respectively (**Fig. 4g**). The number of highly methylated sites in the brain closely correlated with global methylation levels across organs, suggesting that m^6^A deposition may be dictated by global rather than gene-specific regulatory mechanisms. While organ-specific modifications do occur, they appear to be relatively rare and confounded by differences in gene expression. Given that similar trends were observed in both mouse tissues and human cell lines, we believe these largely consistent global methylation patterns reflect biologically relevant regulation rather than artifacts of immortalization or cancer-associated homeostasis.

### m^6^A profiling using limited input

Lastly, we probed the detection limit of eTAM-seq-v2 by lowering the input amount to 100, 20, and 10 ng of total RNA. As poly(A)^+^ selection becomes challenging with limited input^36^, we designed our low-input eTAM-seq-v2 workflow to begin with total RNA. In this workflow, HEK293T total RNA is fragmented, ligated to a biotinylated DNA adapter, and immobilized on streptavidin magnetic beads prior to ribosomal RNA (rRNA) depletion using targeted DNA probes and RNase H (**Fig. 5a**). RNA is then treated with TadA8r and converted into DNA libraries following standard protocols. We observed minimal adapter dimer formation even when reducing the input to 10 ng total RNA (∼500 cells; **Extended Data Fig. 7a**), suggesting that our low-input eTAM-seq-v2 workflow is robust and may support even lower inputs.

**Figure 5.**
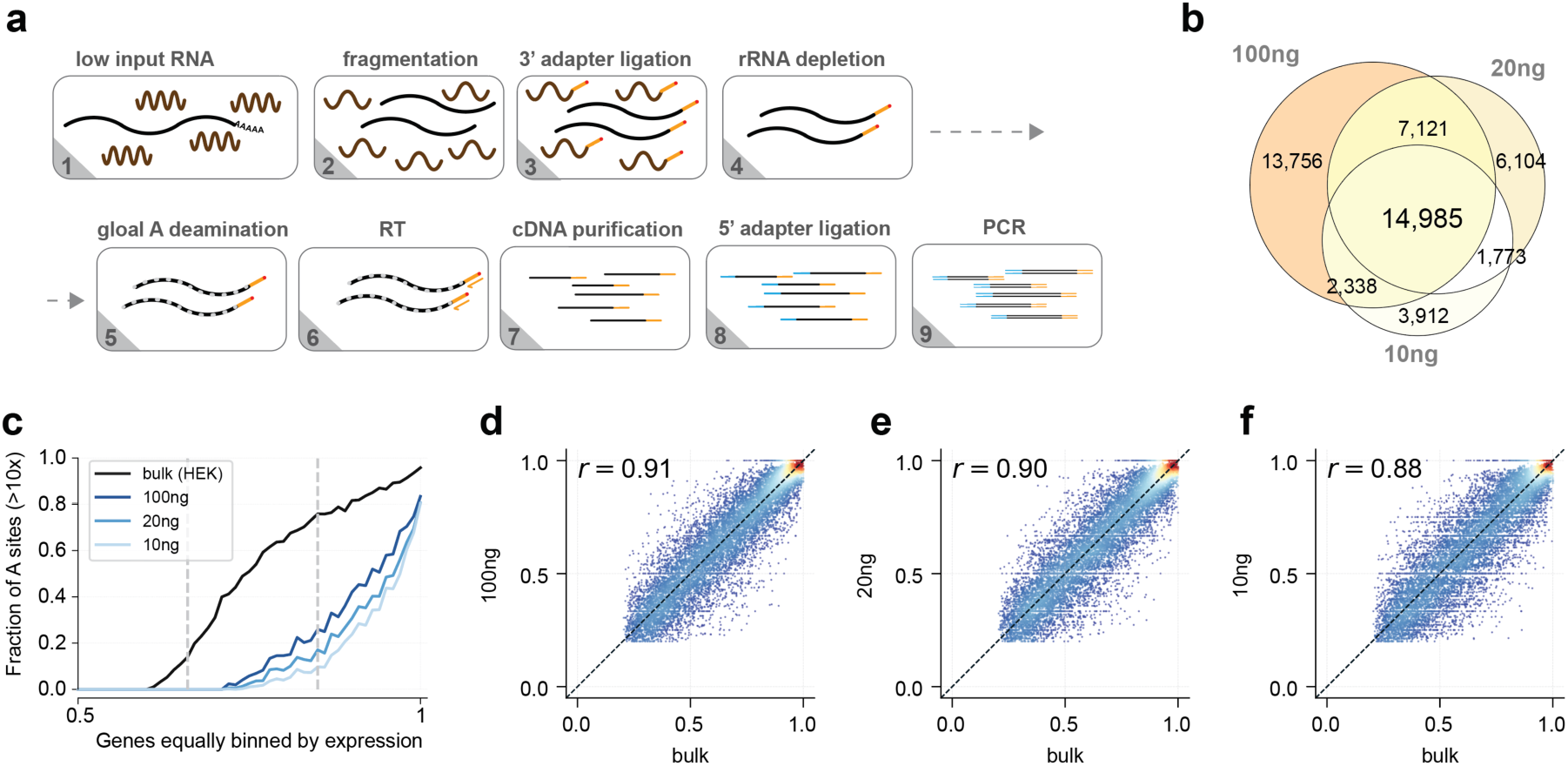
Transcriptome-wide m^6^A profiling with limited input. **a**. Schematic of low-input eTAM-seq-v2. **b**. Venn diagram showing the overlap of m^6^A sites detected using 100, 20, and 10 ng of HEK293T total RNA. **c**. Fraction of effective A sites relative to total A sites in genes binned by expression percentile using 50 ng mRNA, and 100, 20, and 10 ng total RNA. The fraction of effective A sites was calculated as the number of effective A sites detected in each gene divided by the total A sites in the gene. Vertical dashed lines indicate RPKM = 1 and 10. For genes with RPKM >1, 49%, 41%, and 35% of effective A sites detected in bulk samples were recovered. **d**-**f**. Methylation level correlation of m^6^A sites detected in bulk samples and 100 (**d**), 20 (**e**), and 10 ng (**f**) of total RNA. m^6^A sites co-identified in two eTAM-seq-v2 replicates with 50 ng HEK293T mRNA were used to represent bulk results.

We detected 38,200, 29,983, and 23,008 m^6^A sites using 100, 20, and 10 ng total RNA, respectively, with 14,985 sites common to all three samples (**Fig. 5b** and **Supplementary Table 4**). These numbers are lower than the number of m^6^A sites detected in bulk HEK293T mRNA (**Fig. 3b**). To probe the cause of this seemingly reduced sensitivity, we examined the number of effective A sites—those with sufficient sequencing coverage for m^6^A calling—in genes binned by expression levels. The proportion of effective A sites relative to total A sites was lower in low-input samples than in bulk mRNA (35-49% of those detected in bulk samples for genes with RPKM >1; **Fig. 5c**). Therefore, the decrease in detected m^6^A sites reflects reduced sequencing coverage rather than technical bias. Note that fewer reads were allocated to these low-input samples compared to bulk mRNA samples (**Supplementary Table 1**), and we expect coverage to increase with deeper sequencing.

The identified m^6^A sites were enriched in RAC motifs (79-83%) and showed metagene profiles consistent with those observed for bulk samples (**Fig. 3b** and **Extended Data Fig. 7b-d**). Similarly, methylation levels determined using limited input correlated well with those obtained from bulk samples (Pearson’s *r* = 0.88-0.91; **Fig. 5d-f**). In all three samples, we observed minimal reads corresponding to human rRNA (0.2-0.4%; **Supplementary Table 1**) and *E. coli*-derived sequences (0.2-0.4%; likely introduced by TadA8r and other enzymes used during library construction). Collectively, eTAM-seq-v2 supports transcriptome-wide m^6^A detection and quantification with as few as 500 cells.

## Discussion

In this study, we present eTAM-seq-v2, a second-generation eTAM-seq technology that advances m^6^A detection through improved fidelity and sensitivity. We optimized several key parameters during its development. First, we replaced TadA8.20 with TadA8r, which increased the global deamination rate from 91.7% (in eTAM-seq-v1) to 98.6% (in eTAM-seq-v2). This substantial improvement enables direct m^6^A calling without relying on control transcriptomes. Second, we streamlined the workflow by integrating enzymatic treatment into a commercial library preparation kit that uses template switching rather than ligation for adapter installation. This modification standardizes the protocol and reduces processing time from three days to eight hours. Finally, we developed low-input eTAM-seq-v2, which supports transcriptome-wide m^6^A profiling with as little as 10 ng of total RNA.

With eTAM-seq-v2, we delineated the m^6^A landscape in human and mouse mRNA. We found that methylation patterns among six human cell lines and across seven embryonic mouse organs were largely consistent, with global upregulation and downregulation correlating with the expression of the entire m^6^A writer complex. Our observations are corroborated by orthogonal detection methods, as the m^6^A profile reported by eTAM-seq-v2 in HeLa cells is consistent with that mapped by GLORI in HEK cells (**Fig. 2h**). Although differential methylation does occur and is sufficient to separate cell and tissue types, it is considerably rarer than differential expression. This observation is perhaps unsurprising, as expression levels can vary by orders of magnitude across cell types, while methylation has a narrower dynamic range (0-100%). Significant changes in methylation are thus more challenging to arise biologically and be detected technically. Additionally, methylation can only be quantitatively compared among A sites effectively covered in both samples, a parameter inherently compounded with expression. Although a typical eTAM-seq experiment can survey more than half of A sites in expressed genes (RPKM >1), we cannot exclude the possibility that differentially methylated sites tend to reside in lowly expressed genes. Targeted eTAM-seq analysis via amplicon sequencing may help capture these events but remains difficult to scale. Moreover, our analysis is limited to mRNA and does not address differential methylation in other RNA species.

While highly methylated neighboring m^6^A sites must co-occur in the same transcript, an intriguing open question in m^6^A research is whether moderate and low methylation signals are randomly distributed across all RNA molecules transcribed from a given gene or arise from a subset of them. m^6^A-mediated regulation could either be uniformly applied to all transcripts or selectively impact a subset. With eTAM-seq-v2, we demonstrate that most neighboring m^6^A sites in human cells are deposited independently of one another, supporting the notion that m^6^A-mediated regulation does not differ significantly at the single-molecule level. A similar conclusion was recently drawn from GLORI-based analyses in planarians^37^.

In the current study, we focused on m^6^A sites with methylation levels above 20%. While eTAM-seq-v2 is capable of detecting methylation below this threshold (**Extended Data Fig. 2f**), such measurements are inherently noisier. Researchers are therefore encouraged to include appropriate controls to validate low-level methylation events. While TadA8r is highly efficient, it retains a minor sequence bias, particularly at C<u>A</u>G and G<u>A</u>G motifs, which can affect m^6^A calling. As conversion resistance correlates with local GC content, we successfully modeled the background to minimize its impact. An evolved TadA variant lacking sequence bias, or a cocktail of deaminases covering all 16 N<u>A</u>N contexts, could further lower the background and improve eTAM-seq performance. Additionally, increasing the thermostability of TadA8r through protein design could facilitate higher-temperature treatments to resolve RNA secondary structure. Improved folding and stability would also simplify protein overexpression and purification, making the enzyme more accessible.

In summary, eTAM-seq-v2, with its mild enzymatic processing and superior deamination efficiency, enables faithful m^6^A detection from limited RNA input. We envision that eTAM-seq will become a routine tool for researchers to survey m^6^A in diverse biological contexts and continue to drive advances in our molecular understanding of this epitranscriptomic mark.

## Acknowledgements

This work was completed in part with computing resources provided by the University of Chicago Research Computing Center. We thank the Genomics Core and the Single Cell Immunophenotyping Core Facility at the University of Chicago for sequencing support. **Funding**: W.T. is supported by the Searle Scholars Program (SSP-2021-113), the Cancer Research Foundation Young Investigator Program, the American Cancer Society (RSG-22-043-01-ET), the David & Lucile Packard Foundation (2022-74685), and the Sloan Foundation. C.H. is an investigator of the Howard Hughes Medical Institute.

## Author contributions

E.M.H., Y.W., C.Y., and W.T. conceived and designed the study. Y.W. biochemically characterized TadA8r. E.M.H. and Y.W. purified TadA8.20, TadA8r, designed and conducted eTAM-seq-v2. C.Y. analyzed the data with input from E.M.H. and Y.W. T.B.Z. harvested mouse embryos. C.H. and W.T. supervised the study. E.M.H., Y.W., C.Y., and W.T. wrote the manuscript with input from T.B.Z. and C.H.

## Competing interests

The University of Chicago has filed a patent on eTAM-seq. C.H. is a scientific founder, a member of the scientific advisory board, and equity holder of Aferna Bio, Inc. and Ellis Bio Inc., a scientific cofounder and equity holder of Accent Therapeutics, Inc., and a member of the scientific advisory board of Rona Therapeutics and Element Biosciences.

## Data availability

All sequencing data have been deposited to the Gene Expression Omnibus (GEO) under accession number GSE294952.

## Code availability

Codes for analyzing eTAM-seq-v2 libraries are available in the following GitHub repository: https://github.com/y9c/m6A-eTAMseq

## Methods

### General methods

All oligonucleotides were synthesized by Integrated DNA Technologies (IDT). DNA was amplified using Taq DNA Polymerase (New England BioLabs, catalog no. M0273X) or NEBNext Ultra II Q5 Master Mix (NEB, cat. no. M0544X). DNA was purified using the QIAquick PCR Purification Kit (Qiagen) or the DNA Clean & Concentrator-5 Kit (Zymo research). RNA was purified using the RNA Clean & Concentrator-5 Kit (Zymo research). Chemicals were purchased from Fisher Scientific or Sigma-Aldrich and used without purification unless otherwise noted. Key plasmids for reproducing this work have been deposited to Addgene (194702 and 208318).

### Cell culture

Cell lines were purchased from American Type Culture Collection. HeLa, HEK293T, and HepG2 cells were maintained in Dulbecco’s Modified Eagle’s Medium (DMEM) supplemented with 10% fetal bovine serum (FBS) at 37°C under 5% CO_2_. U87, Jurkat, and K562 cells were maintained in Roswell Park Memorial Institute (RPMI) 1640 medium supplemented with 10% FBS at 37°C under 5% CO_2_.

### Mouse embryos

Mice were housed in a specific pathogen-free facility with a 12-h light-dark cycle. C57BL/6J mice were purchased from the Jackson Laboratory (JAX# 000664) and bred in house. Male and female adult mice were used to set up timed mating. The presence of a vaginal plug in the morning was defined as embryonic day 0.5 (E0.5). Mouse tissues were dissected in PBS from embryos at E18.5, flash frozen on dry ice, and stored at – 80 °C. All animal experiments were approved by the Institutional Animal Care and Use Committee at the University of Chicago under protocol 72541.

### Overexpression and purification of recombinant TadA8.20 and TadA8r protein

TadA8r and TadA8.20 were overexpressed and purified following previously reported protocols^9, 14^. Briefly, TadA8.20 or TadA8r were cloned into a pET28a vector following an N-terminal, hexahistidine-tagged maltose-binding protein (6×His-MBP). A tobacco etch virus (TEV) protease cleavage site (ENLYFQ|G) was installed between MBP and TadA for post-expression tag removal. BL21 Rosetta (DE3) competent cells were transformed with the expression plasmids and plated on Luria broth (LB) agar. Unless otherwise noted, antibiotics were supplied at fixed concentrations: 50 μg mL^−1^ for kanamycin (for expression plasmid maintenance) and 32 μg mL^−1^ for chloramphenicol (for BL21 Rosetta). Single colonies were inoculated into fresh LB medium and grown in an incubator shaker (37 °C, 220 r.p.m.) until the absorbance at 600 nm (OD_600_) reached ∼0.8. This culture was then inoculated into fresh LB (20 mL per 1 L) for protein overexpression. The culture was grown to OD_600_ ∼0.6, cooled immediately to 4 °C, and induced with 0.1 mM isopropyl β-d-1-thiogalactopyranoside (IPTG). Bacteria were cultured at 16 °C for an additional 20 h before pelleting by centrifugation at 4,000 *g*.

Bacteria were lysed by sonication in buffer A (50 mM Tris, 500 mM NaCl, 10 mM 2-mercaptoethanol and 10% (v:v) glycerol, pH 7.5). Lysed bacteria were clarified by centrifugation at 4 °C and 23,000 *g*. The supernatant was loaded onto a Ni-NTA Superflow Cartridge (QIAGEN, cat. no. 30761), washed with 30 mL of buffer A supplemented with 50 mM imidazole, and eluted with a gradient of imidazole from 50 mM to 500 mM in buffer A. The eluted protein was buffer exchanged to remove imidazole and dialyzed with TEV protease (∼3 mM in a 5 mL dialysis cassette) in buffer B (50 mM Tris, 200 mM NaCl, 10 mM 2-mercaptoethanol and 10% (v:v) glycerol, pH 7.5) at 4 °C overnight.

For TadA8.20, the protein mixture was reloaded onto a Ni-NTA Superflow Cartridge, washed with buffer C (50 mM Tris, 1 M NaCl, 10 mM 2-mercaptoethanol and 10% (v:v) glycerol, pH 8.0), and eluted by buffer C supplemented with 50 mM imidazole. For TadA8r, the protein mixture was loaded onto a UNOsphere S Column (BioRad, cat. no. 12009305), washed with buffer B (50 mM Tris, 200 mM NaCl, 10 mM 2-mercaptoethanol and 10% (v:v) glycerol, pH 7.5), and eluted with a NaCl gradient from 200 mM to 1 M in buffer B.

Finally, tag-free TadA8.20 or TadA8r was purified by size-exclusion chromatography (Enrich SEC 650 10 × 300 mm^2^ column; BioRad, cat. no. 7801650). The column was balanced and eluted with buffer B. The resulting protein was buffer exchanged and concentrated using a 10 kDa MWCO filter (Cytiva, cat. no. 28932296) into approximately 4 mg mL^−1^ in buffer B (prepared with molecular grade water) before being flash frozen for future use. Starting from a 6 L culture, we typically get 200 μL purified TadA8.20 or TadA8r.

### LC Q-TOF MS quantification of A-to-I conversion in an RNA probe

An RNA probe (UUUACUUU; 5 ng) was incubated with 7 μL of deaminase (diluted to different concentrations in buffer B) in 30 μL 1× deamination buffer (50 mM Tris, 25 mM KCl, 2.5 mM MgCl_2_, 2 mM DTT and 10% (v:v) glycerol, pH 7.5). The reaction was incubated at 44 °C for 3 h. Proteinase K (Fisher Scientific, cat. no. BP1700) was added to a final concentration of 200 ng μL^−1^, incubated at 37 °C for 10 min, 55 °C for 1 h followed by deactivation at 85 °C for 30 min and 95 °C for 10 min. The sample was clarified by centrifugation at 21,300× *g* for 10 min, and 20 μL of the supernatant was analyzed by LC Q-TOF MS.

High-performance liquid chromatography was performed using a reverse-phase C18 column (Agilent, Poroshell 120 EC-C18, 3.0 × 150 mm, 2.7 μm, PN 695575-302). Fractionated samples were directly analyzed using an Agilent 6540 UHD Q-TOF LC mass spectrometer in a negative electrospray ionized mode (AJS-ESI, Gas Temp 300 °C, Gas Flow 13 L/min, Nebulizer 35 psig, SheathGasTemp 350 °C, SheathGasFlow 12 L/min, Fragmentor Voltage 190V). The solvent system used for LC was: solvent A = 25 mM 1,1,1,3,3,3-hexafluoro-2-propanol (AA blocks, cat. no. 920-66-1) and 15 mM hexylamine (Sigma-Aldrich, cat. no. 219703) in water; solvent B = methanol. The solvent program was set as: 20% B for 2 min; 20-60% B over 8 min; 60% B for 2 min; 60-100% B over 1 min; 100% B for 2 min.

Unconverted and converted RNA was quantified using Agilent MassHunter Qualitative Analysis 10.0 in an extracted ion chromatogram (EIC) mode. Data were fitted using a nonlinear regression model in GraphPad.

### *In vitro* deamination of DNA by TadA8r and TadA8.20

A DNA probe (GTGGTGCGGCCTGCTGTGGCCCTCATGGACATATGTTGGTGTTGGCTGGGTTTGGGGGTC; 20 nM) was incubated with 10/1/0.1 μM TadA8r or TadA8.20 in 20 μL 1× deamination buffer (50 mM Tris, 25 mM KCl, 2.5 mM MgCl_2_, 2 mM DTT and 10% (v:v) glycerol, pH 7.5) at 37 °C for 1 h. Reactions were quenched by incubating at 95 °C for 10 min. PCR was set up with 1 μL of reaction mixture as the template, and followed the program with Taq polymerase: denaturing (95 °C for 3 min); 25 cycles of amplification (denaturing at 95 °C for 10 s, annealing at 60 °C for 10 s followed by extension at 68 °C for 20 s) and final extension at 68 °C for 5 min. Primers used were PCR_Fwd (TGCGGCCTGCTGTGGC) and PCR_Rev (CAAACCCAGCCAACACCAAC). PCR products were purified and analyzed by Sanger sequencing.

### Melting temperature measurement

TadA8.20 or TadA8r (4 μg) was incubated with 5× SYPRO Orange dye (Invitrogen, cat. no. S6651) in 10 μL deamination buffer (50 mM Tris, 25 mM KCl, 2.5 mM MgCl_2_, 2 mM DTT and 10% (v:v) glycerol, pH 7.5). Samples were heated from 10 °C to 95 °C in increments of 0.5 °C and held for 10 s before measuring fluorescence in a CFX96 Real-time PCR system (BioRad). The fluorescence signal was detected in the FRET channel following a protein thermal shift assay protocol developed by BioRad.

### Heat deactivation of TadA8r

Fluorescein-labeled ssDNA (/56-FAM/TGGGTTGGTT<u>A</u>TCGTTTGGTGG; 1 μM) was pre-incubated in 9 μL 1× deamination buffer (50 mM Tris, 25 mM KCl, 2.5 mM MgCl_2_, 2 mM DTT and 10% (v:v) glycerol, pH 7.5) at specific temperatures for 20 min. 1 μL of 100 μM TadA8r was preheated to matching temperatures for 1 min before being mixed with the DNA substrate. Reactions were incubated at the same temperature for 1 h and quenched by incubating at 95 °C for 10 min. TadA8r was digested by Proteinase K (Fisher Scientific, cat. no. BP1700) at a final concentration of 200 ng μL^−1^ (incubation at 55 °C for 2 h followed by deactivation at 85 °C for 30 min and 95 °C for 15 min). The reaction was then incubated with 10 U of *E. coli* endonuclease V (New England Biolab, cat. no. M0305S) at 37 °C for 2 h. After endonuclease V digestion, a 5 μL aliquot was mixed with 5 μL 2× loading buffer (95% formamide, 10 mM EDTA, 0.025% SDS, and 0.025% (w/v) bromophenol blue), heated to 95 °C for 5 min, and resolved by 15% (v/v) denaturing polyacrylamide gel. ssDNA was visualized in ChemiDoc XRS+ (Bio-Rad) using the fluorescein channel and quantified by ImageJ. Heat deactivation curve was plotted in GraphPad.

### Poly(A)^+^ RNA purification and fragmentation

Cells or tissues were rinsed with 1× PBS and lysed by the direct addition of TRIzol reagent (Invitrogen, cat. no. 15596026). Tissue samples were further homogenized with a homogenizer. Total RNA was then purified following the manufacturer’s protocol, while Poly(A)^+^ RNA was enriched from total RNA using Dynabeads mRNA Purification Kits (Invitrogen, cat. no. 61006). For standard eTAM-seq, 50 ng of poly(A)^+^ RNA was fragmented with a magnesium RNA fragmentation module (NEB, cat. no. E6150S) following the manufacturer’s recommendation. Fragmented RNA was purified by RNA Clean & Concentrator kits (Zymo Research, cat. no. R1014).

### Standard eTAM-seq library preparation

Fragmented RNA was treated with 200 pmol TadA8r in 20 μL deamination buffer (50 mM Tris, 25 mM KCl, 2.5 mM MgCl_2_, 2 mM DTT and 10% (v:v) glycerol, pH 7.5) supplemented with 10% (v:v) SUPERase•In RNase Inhibitor at 53 °C for 1 h, followed by purification using RNA Clean & Concentrator kits. The reaction was repeated two times at 44 °C for 1 h, each time followed by column purification, resulting in 3x reactions in total. TadA8r-treated RNA was then used as the starting material for the SMARTer Stranded Total RNA-Seq Kit v3 (Takara, cat. no. 634485), following the manufacturer’s protocol option 2 (without fragmentation). Typically, 10-11 cycles of PCR were carried out to generate the final library, which was purified by AMPure XP beads (Beckman Coulter, cat. no. A63882) following the manufacturer’s directions and submitted for NGS.

### Low-input eTAM-seq library preparation

HEK293T total RNA (100 ng, 20 ng, or 10 ng) was fragmented using a magnesium RNA fragmentation module (NEB, cat. no. E6150S) following the manufacturer’s recommendation and purified with Oligo Clean & Concentrator kits (Zymo Research, cat. no. D4060). Fragmented RNA was treated with 50 U of T4 polynucleotide kinase (NEB, cat. no. M0201L) in 50 μL T4 PNK Buffer (100 mM NaAc, 10 mM MgCl_2_, 5 mM DTT, 10% (v:v) SUPERase•In RNase Inhibitor, pH 6.0) at 23 °C for 1 h and purified with Oligo Clean & Concentrator kits. End-repaired RNA was ligated to 20 pmol of 3’-adapter (/5rApp/AGATCGGAAGAGCGTCGTG/3Bio/) using 400 U of T4 RNA ligase 2, truncated KQ (NEB, cat. no. M0373L) supplemented with 8% (v:v) SUPERase•In RNase Inhibitor and 15% (w/v) PEG 8000 in a 25 μL reaction at 25 °C for 4 h. The reaction was further incubated at 16 °C overnight. Excess adapters were first digested by 5’-deadenylase (NEB, cat. no. M0331S) at 30 °C for 1 h, followed by RecJ_f_ (NEB, cat. no. M0264L) at 37 °C for 1 h.

Ligated RNA was immobilized onto 10 μL of Dynabeads MyOne Streptavidin C1 (Invitrogen, cat. no. 65002) in 1× Binding/Wash Buffer (5 mM Tris-HCl, 1 M NaCl, 0.5 mM EDTA, pH 7.5). rRNA was depleted with the NEBNext rRNA depletion kit v2 (NEB, cat. no. E7400) following the manufacturer’s protocol. Remaining RNA was then deaminated on beads with 200 pmol TadA8r in 20 μL 1× deamination buffer supplemented with 10% (v:v) SUPERase•In RNase Inhibitor at 53 °C for 1 h. 3.5 μL of beads were supplemented to the reaction mixture, incubated for 10 min, before the beads were decanted, washed once by 1× Binding/Wash Buffer, and twice in a low-salt buffer (10 mM Tris-HCl, 50 mM NaCl, pH 7.5). The deamination reaction was repeated twice at 44 °C for 1 h, each with the replenishment of beads and wash, resulting for 3 treatments in total.

RNA was annealed to 2 pmol of RT primer (ACACGACGCTCTTCCGATCT) at 70 °C for 2 min, and then reverse transcribed with 200 U of Maxima H^−^ Reverse Transcriptase (Thermo Fisher Scientific, cat. no. EP0753) in 25 μL 1× RT buffer, with 5 mM RNaseOUT (Invitrogen, cat. no. 10777019) and 0.5 mM dNTP at 50 °C for 1 h. cDNA was released by boiling the beads in 0.1% (w:v) sodium dodecylsulfate (SDS) at 95 °C for 10 min. The elute was purified by DNA Clean & Concentrator kits (Zymo Research, cat. no. D4014). Purified cDNA was ligated to a UMI-containing cDNA adapter (/5Phos/NNNNNNAGATCGGAAGAGCACACGTCTG/3SpC3/) using 30 U of T4 RNA ligase (NEB, cat. no. M0437M) in 50 μL 1× T4 RNA ligase reaction buffer in the presence of 1 mM ATP, 25% (w/v) PEG 8000, 7.5% (v/v) DMSO and 1 mM CO(NH_3_)_6_Cl_3_. The ligation reaction was incubated at 25 °C for at least 14 h before purification using DNA Clean & Concentrator kits. Adapter-containing cDNA was then PCR amplified with the NEBNext Ultra II Q5 Master Mix (NEB, cat. no. M0544X) and NEBNext Unique Dual Index Primers for Illumina (NEB, cat. no. E6440S) following the manufacturer’s directions. The resulting DNA library was purified by AMPure XP beads (Beckman Coulter, cat. no. A63882) following the manufacturer’s directions and submitted for NGS.

### Overexpression and purification of FTO

FTO was overexpressed and purified following previously reported methods^9, 38^. Briefly, the human FTO gene was cloned into a pET28a vector and transformed into BL21 Rosetta (DE3). Successfully transformed bacteria were used to start an overnight culture, which was subsequently diluted into fresh 2× YT broth (1:50) and cultured at 37 °C to an OD_600_ of 0.8–1.0. The culture was cooled to 16 °C and supplemented with 0.1 mM IPTG, 10 μM ZnSO_4_ (Sigma-Aldrich), and 2 μM (NH_4_)_2_Fe(SO_4_)_2_ (Sigma-Aldrich). Bacteria were cultured overnight at 16 °C before collection via centrifugation.

Bacteria were lysed in buffer D (300 mM NaCl, 50 mM imidazole, and 50 mM of Na_2_HPO_4_, pH 8.0). The lysate was clarified by centrifugation, loaded onto a nickel column, washed with buffer E (150 mM NaCl, 25 mM imidazole, and 10 mM Tris-HCl, pH 7.5), and eluted with buffer F (150 mM NaCl, 250 mM imidazole, and 10 mM Tris-HCl, pH 7.5). The eluate was loaded on to an anion-exchange column (SOURCE 15Q, Cytiva) and fractionated with 0–50% buffer G (1.5 M NaCl, 20 mM Tris-HCl, pH 7.5) over 30 min. The resulting protein was buffer exchanged and concentrated using a 10 kDa MWCO filter (Cytiva, cat. no. 28932296) before being flash frozen in 30% glycerol for future use.

### Preparation of FTO-treated transcriptome samples

Fragmented RNA was demethylated by incubating with 200 pmol of FTO in 20 μL 1× FTO reaction buffer (2 mM sodium ascorbate (Sigma-Aldrich), 65 μM (NH_4_)_2_Fe(SO_4_)_2_, 0.3 mM α-ketoglutarate (Sigma-Aldrich), 0.1 mg mL^−1^ bovine serum albumin (NEB), and 50 mM HEPES-KOH, pH 7.0) supplemented with 10% (v:v) SUPERase•In RNase Inhibitor (Invitrogen, cat. no. AM2696) at 37 °C for 1 h, and then purified by RNA Clean & Concentrator kits.

### Preparation of *in vitro* transcribed transcriptomes

Modification-free control transcriptomes were prepared from various mRNA samples based on previously published protocols^9^. An oligo-dT(30)VN primer (TTTTTTTTTTTTTTTTTTTTTTTTTTTTTTVN, 100 pmol) was annealed to 100 ng of poly(A)^+^ RNA at 65 °C for 5 min. RNA was then reverse transcribed in 20 μL 1× RT buffer (Thermo Fisher Scientific, cat. no. EP0753; 50 mM Tris-HCl, 75 mM KCl, 3 mM MgCl_2_ and 10 mM DTT, pH 8.3) in the presence of 40 pmol of 5Bio-T7-TSO (/5Biosg/ACTCTAATACGACTCACTATAGGGAGAGGGCrGrGrG), 1 mM of GTP, 5% (w:v) PEG 8000, 0.5 mM of each dNTP, 5 mM RNaseOUT (Invitrogen, cat. no. 10777019) and 200 U of Maxima H− Reverse Transcriptase (Thermo Fisher Scientific, cat. no. EP0753). The reaction proceeded under the following condition: 42 °C for 90 min, 10 cycles of 50 °C for 2 min + 42 °C for 2 min, and 85 °C for 5 min. Subsequently, 10 μL of RNase H (NEB, cat. no. M0297L), 70 μL of RNase-free H_2_O, and 100 μL of Ultra II Q5 Master Mix (NEB, cat. no. M0544X) were added to make the second-strand synthesis mixture, which was incubated under the following condition: 37 °C for 15 min, 95 °C for 1 min, and 65 °C for 10 min. The reaction was purified with 160 μL (0.8×, v:v) of AMPure XP beads (Beckman Coulter, cat. no. A63882) following the manufacturer’s directions.

Purified and concentrated dsDNA was *in vitro* transcribed in 20 μL 1× T7 Reaction Buffer (NEB, cat. no. E2040S; 40 mM Tris-HCl, 6 mM MgCl_2_, 1 mM DTT, and 2 mM spermidine, pH 7.9) with 10 mM of each NTP and 2 μL of T7 RNA Polymerase Mix (NEB, cat. no. E2040S) at 37 °C overnight. The IVT mixture was further treated with TURBO DNase (Invitrogen, cat. no. AM2238) and purified by acid–phenol chloroform (Invitrogen, cat. no. AM9722) extraction and ethanol precipitation to yield 2.5–10 μg of IVT RNA. The IVT transcriptome was then subjected to the standard library preparation protocol.

### Sequencing data processing

Adapter sequences and low-quality bases (Phred score <20) at the 3’ end were trimmed from the raw reads using Cutadapt^39^. Reads shorter than 20-nt were subsequently removed. For standard eTAM-seq samples prepared using the random primming method, the 8-nt sequences at the 5’ end of the R2 reads were extracted to obtain unique molecular identifiers (UMIs). Following the SMARTer Stranded Total RNA-Seq Kit v3 protocol, 3-nt from the 5’ end of R1 and 6-nt from the 5’ end of R2 were removed. For eTAM-seq low-input samples prepared using the ligation method, the 6-nt sequences at the 5’ end of the R2 reads were extracted to obtain UMI. Subsequently, 2-nt from the 5’ end of R1 and 1-nt from the 5’ end of R2 were removed.

For head-to-head comparison of eTAM-seq and GLORI, three GLORI libraries, HEK293T ligation-based (SRR21356251), HeLa ligation-based (SRR21356219), and HEK293T random priming-based (SRR21356199) were downloaded from the NCBI database. Similar adapter trimming strategies were applied to ligation-based and random primming-based libraries, respectively, except that for ligation-based libraries, the 10-nt sequence at the 5’ end of R2 was extracted as the UMI.

rRNA and tRNA genes (human or mouse) and *E. coli* sequences were initially masked using HISAT-3N^40^ with following parameters: “--base-change A,G --no-spliced-alignment --no-softclip --directional-mapping”. The remaining reads were mapped to the reference genome (GRCh38 for human cell line samples, GRCm39 for E18.5 mouse embryo samples) using HISAT-3N (--base-change A,G --bowtie2-dp 1 –score-min L,0,1 -- directional-mapping). Deduplication was done by the subcommand dedup from UMI-tools using the UMI extracted prior to alignment. To minimize mapping errors, reads with >5% of mismatches (excluding A-to-G conversions) were discarded. To further mitigate potential impact from non-templated nucleotides sometimes added during RT, the first and last 2 nucleotides of each read were masked before site detection.

### Library quality assessment

The A-to-G global deamination rate was calculated by dividing G counts by the combined A and G counts mapped to genomic A sites. The C-to-T conversion rate was calculated by dividing T counts by the combined C and T counts mapped to genomic C sites. The insert size was calculated as the distance between the 5’ ends of R1 and R2 in each paired read. For GLORI libraries, which distributed only R2 reads, the insert size was estimated based on the mapping length. This approach is valid because the sequencing read length (150 nt) is substantially longer than the average insert size (∼40 nt).

### Coverage assessment

To assess coverage across the four nucleotides (A/T/G/C), site coverage within individual genes was internally normalized by the mean coverage per gene. This normalization ensured that each gene contributed equally, preventing highly expressed genes from disproportionately influencing the results. The normalized coverages across all genes were then averaged for each base. Finally, the coverage for each base was presented as the relative coverage of A, T, G, and C. In the rarefaction analysis, A sites covered by more than 10 reads are defined as effective A sites and plotted as a function of the number of uniquely mapped reads. Subsampling was performed based on the observed coverage profile of all sites across the genome, with 10%, 20%, …, up to 100% of the total coverage randomly subsampled, weighted by the observed coverage of each site. This generated a new coverage profile for each site, and sites covered by more than 10 reads were counted.

### Gene expression level quantification

Gene expression levels for all libraries, encompassing 12 cell line samples, 14 mouse tissue samples, and 3 low-input samples, were quantified from eTAM-seq reads using featureCount. This quantification was performed against 38,549 protein-coding and lncRNA genes annotated by ENCODE. The resulting counts were converted to the Reads Per Kilobase per Million mapped reads (RPKM) metric. Based on this metric, genes were categorized as lowly expressed (RPKM >1) or moderately expressed (RPKM >10). Additionally, the RPKM values for known m^6^A writer, eraser, and reader proteins were extracted for each corresponding sample for expression analysis.

### Detection of m^6^A sites

To mask poorly processed RNA fragments, reads with 3 or more consecutive unconverted As were discarded. The remaining reads were used to calculate persistent A rates for individual A sites—total reads of A divided by the total reads of A+G—and therefore methylation levels. GC content of a given A was determined in a 21-nt window including 10 nt upstream and 10 nt downstream sequences flanking the target A. The definition and application of the GC filter are detailed in **Supplementary note 1**. *P* value was calculated for each candidate site. Sites with sequencing depth >10, persistent A rate ≥0.2, and a *P*-value <0.0001 were extracted as m^6^A sites. Methylation levels are determined as persistent A rates.

### Overlapping and correlation between m^6^A sites

For biological replicates, sites that are identified as m^6^A in at least one replicate and covered by more than 10 reads in both replicates were included in the analysis. Pearson correlation coefficients were calculated based on m^6^A modification levels.

To compare two different biological samples (e.g., two distinct cell lines), coverage for each biological sample was calculated as the sum of all its replicates, while m^6^A levels were calculated as the average across replicates. Sites that are identified as m^6^A in any replicate and exhibited coverage greater than 10 in both biological samples were included in the analysis.

For comparisons with eTAM-seq-v1 and GLORI, site lists were directly downloaded from the original studies. Sites with a reported modification level ≥0.2 were included in the analysis.

### Analysis of m^6^A characteristics relative to gene expression

Genes were ranked by expression levels (RPKM) and categorized into 100 equally sized expression bins. To assess gene availability for m^6^A detection relative to expression, the number of genes possessing at least one or at least 100 effective A sites (sites covered by >10 reads for potential m^6^A detection) within exonic regions was determined for each bin. Additionally, the number of genes containing at least one m^6^A site was enumerated. To assess site availability, the average number of total A sites and effective A sites per gene was calculated for each expression bin. The ratio of effective A sites to total A sites was regarded as the total site availability metric.

m^6^A characteristics across expression levels were evaluated for each expression bin by calculating the following metrics: average methylation ratio, defined as the sum of methylation levels divided by the total number of detected m^6^A sites; m^6^A site fraction, defined as the number of detected m^6^A sites divided by the total number of effective A sites; methylation load, defined as the sum of methylation levels of individual sites divided by the total number of effective A sites.

### Co-occurrence analysis

m^6^A sites, as previously defined, with a modification level ≥20%, sequencing depth >10, *P*-value <10^-4^, and located within exonic regions were included for co-occurrence analysis. For each gene, all m^6^A pairs were exhaustively examined, with genes of no more than one m^6^A site excluded. The distance between two sites was calculated based on spliced transcripts, excluding introns. R1 and R2 reads from the same fragment were treated as a single unit, and only fragments simultaneously covering both candidate m^6^A sites (s1 and s2) were used. Modification frequencies at s1 and s2 were counted using all spanning reads and summarized into a contingency table (see **Extended Data Fig. 4a**). Under the null hypothesis (*H_0_*) that the installation of one m^6^A site is independent of the other, *P*-values were calculated using Fisher’s exact test. m^6^A pairs with *P*-values >0.01 were considered independent.

### Differential expression analysis

Differential expression analysis between sample pairs (e.g., mouse brain vs. other organs, or Jurkat vs. other cell lines) was performed using the DESeq2 package^41^. Read counts corresponding to ENCODE-annotated genes, derived by averaging biological replicates, served as input. Genes were identified as differentially expressed if they exhibited at least a two-fold change in expression levels between the compared conditions and had a DESeq2 Wald test adjusted *P*-value less than 0.01.

### Differential methylation analysis

Differential methylation analysis was conducted by comparing m^6^A methylation levels at individual A sites between sample pairs. For pairwise comparison, only sites identified as m^6^A in at least one sample (6 human cell lines, or 7 mouse tissues) and possessing an aggregated sequencing depth greater than 20 (combining reads from biological replicates) in both samples were considered. A site was defined as differentially methylated if it exhibited at least a two-fold difference in methylation levels between the two conditions. For visualization purposes only, log_2_ fold changes of methylation levels were calculated after adding a minimal methylation level of 0.005 to the average methylation level. To assess statistical significance, paired t-tests were performed on the methylation levels measured across the biological replicates, with a significance threshold of *P* < 0.01.

---

**Extended Data Figure 1.**
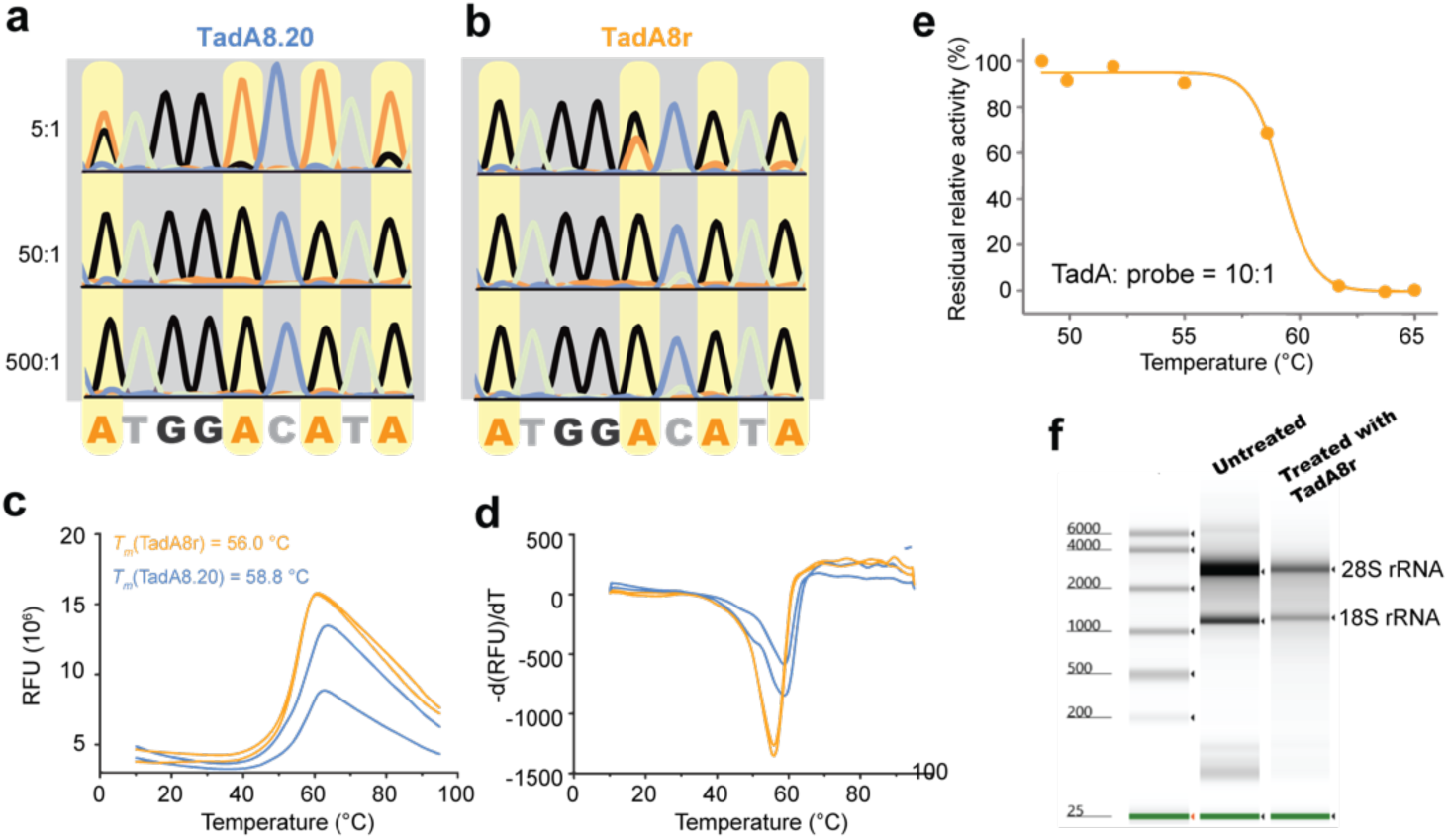
Performance comparison between TadA8.20 and TadA8r. **a**-**b.** TadA8.20- and TadA8r-mediated deoxyadenosine deamination. A DNA probe (20 nM) was incubated with various concentrations of TadA8.20 or TadA8r for 1 h before being amplified by PCR and analyzed by Sanger sequencing. **c**-**d.** Melting curve of TadA8r (orange) and TadA8.20 (blue) quantified with the SYPRO fluorescent dye in TadA reaction buffer. **e.** TadA8r-mediated DNA deamination at different incubation temperatures. ssDNA (1 μM) was incubated with 10 μM TadA8r at different temperatures. Reactions were quenched after 1 h. DNA was purified, processed by Endonuclease V, which cleaves at the 3’ end of the nucleotide following deoxyinosine, and analyzed by polyacrylamide gel electrophoresis (PAGE). Bands corresponding to the starting material and deamination product were quantified using ImageJ. Data were fitted using a nonlinear regression model in GraphPad. *n* = 2 independent experiments. **f.** HeLa total RNA treated with TadA8r at 53 °C for 1 h in reaction buffer (50 mM Tris, 2 mM DTT, 2 mM EDTA, 0.2 mg/mL BSA, 10% (v/v) glycerol, pH = 7.5). EDTA was included to minimize RNA degradation mediated by divalent cations.

**Extended Data Figure 2.**
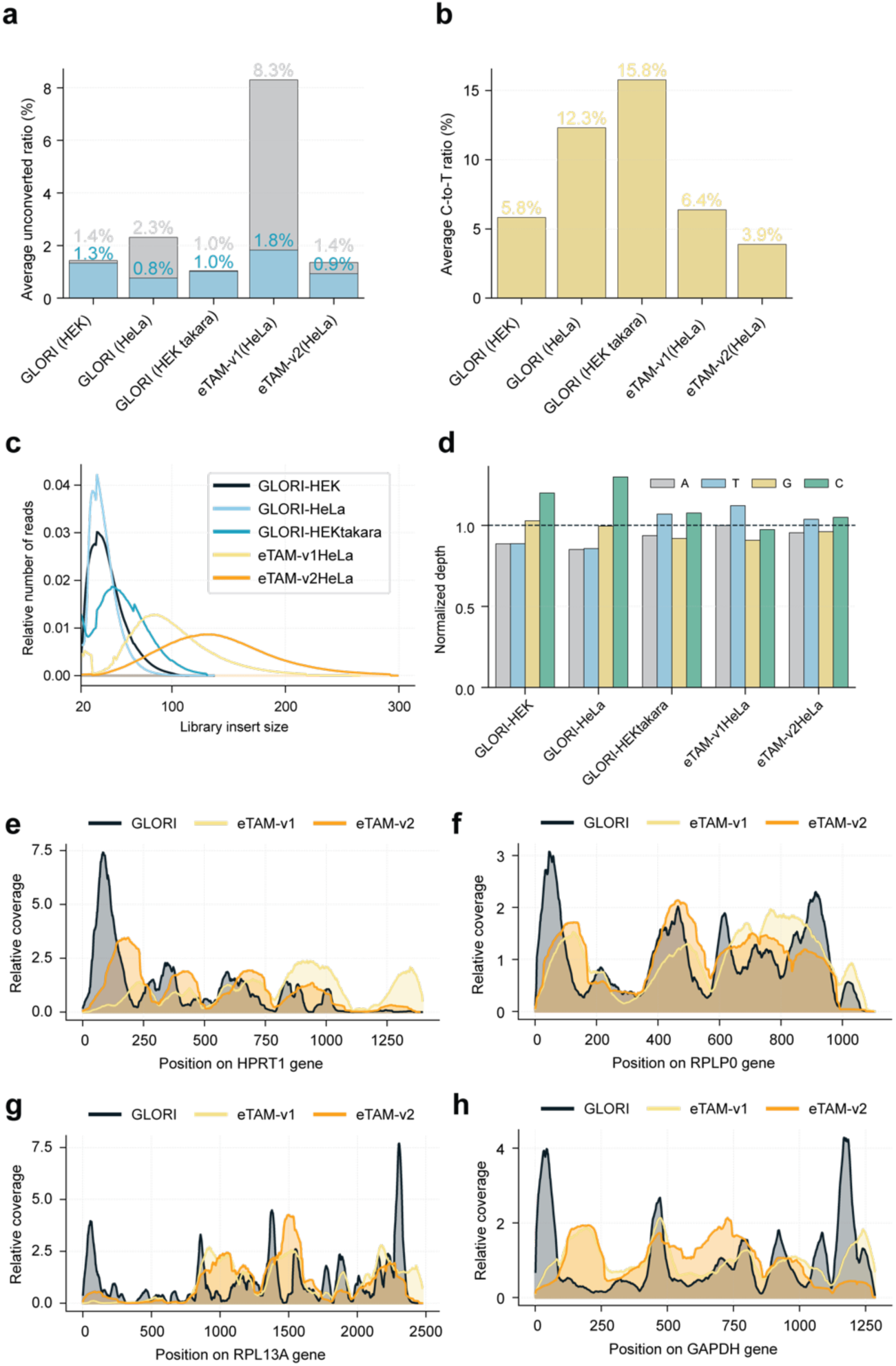
Comparison of sequencing metrics among eTAM-seq-v2, eTAM-seq-v1, and GLORI. GLORI data (HEK293T ligation-based: SRR21356251; HeLa ligation-based: SRR21356219; HEK293T random priming-based: SRR21356199) were obtained from the NCBI SRA and reanalyzed using the same pipeline as the eTAM-seq method. **a.** Average unconverted A rates before (grey) and after (blue) filtering reads with ≥3 consecutive unconverted As. **b.** Average C-to-T mutation rates, calculated from uniquely mapped reads without A conversion filtering. **c.** Distribution of insert lengths. **d.** Normalized base composition. **e-h.** Relative coverage across selected housekeeping genes (*HPRT1*, *RPLP0*, *RPL13A*, and *GAPDH*) with eTAM-seq-v2, eTAM-seq-v1, and GLORI (using the HEK293T ligation-based sample, SRR21356251).

**Extended Data Figure 3.**
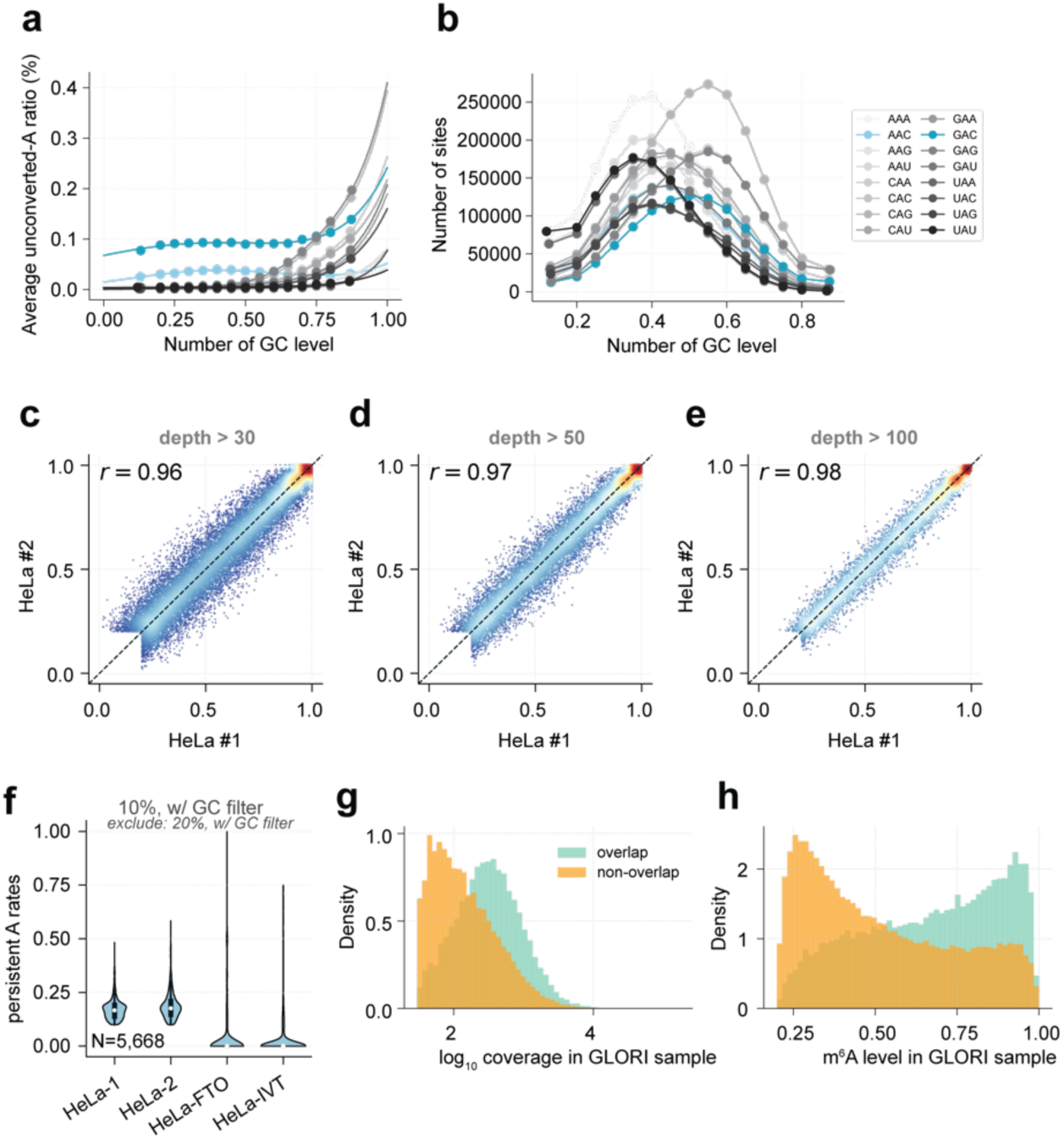
Background analysis and methylation quantification. **a**. Relationship between adenosine conversion resistance and local GC content. A sites with sufficient coverage (depth >10) were binned by their GC content within a 20-nucleotide (nt) window (excluding the central A site). Bins with GC content >80% or <20% were merged, respectively, due to an insufficient number of data points. The average conversion resistance is plotted and fitted as a function of GC content (see **Supplementary Note I**). **b.** Number of A sites analyzed within each bin defined in **a**. **c**-**e.** Correlation of methylation levels between eTAM-seq-v2 replicates using different depth cutoffs (30x, 50x, and 100x). **f.** Violin plot of methylation levels detected in two eTAM-seq-v2 replicates, FTO-treated HeLa mRNA, and an IVT HeLa transcriptome. m^6^A sites with 1) ³10% methylation in both eTAM-seq-v2 replicates; 2) <20% methylation in at least one replicate; 3) sequencing depth >1 in both FTO and IVT samples are plotted. Median is denoted with a dot. **g**-**h.** Comparison of sequencing depth and methylation levels between m^6^A sites co-detected by GLORI and eTAM-seq-v2 (green) and those detected only by GLORI (orange). Due to its higher sequencing depth, the GLORI dataset identified more m^6^A sites overall. GLORI-only sites generally have lower depth and methylation levels than those co-detected by both methods.

**Extended Data Figure 4.**
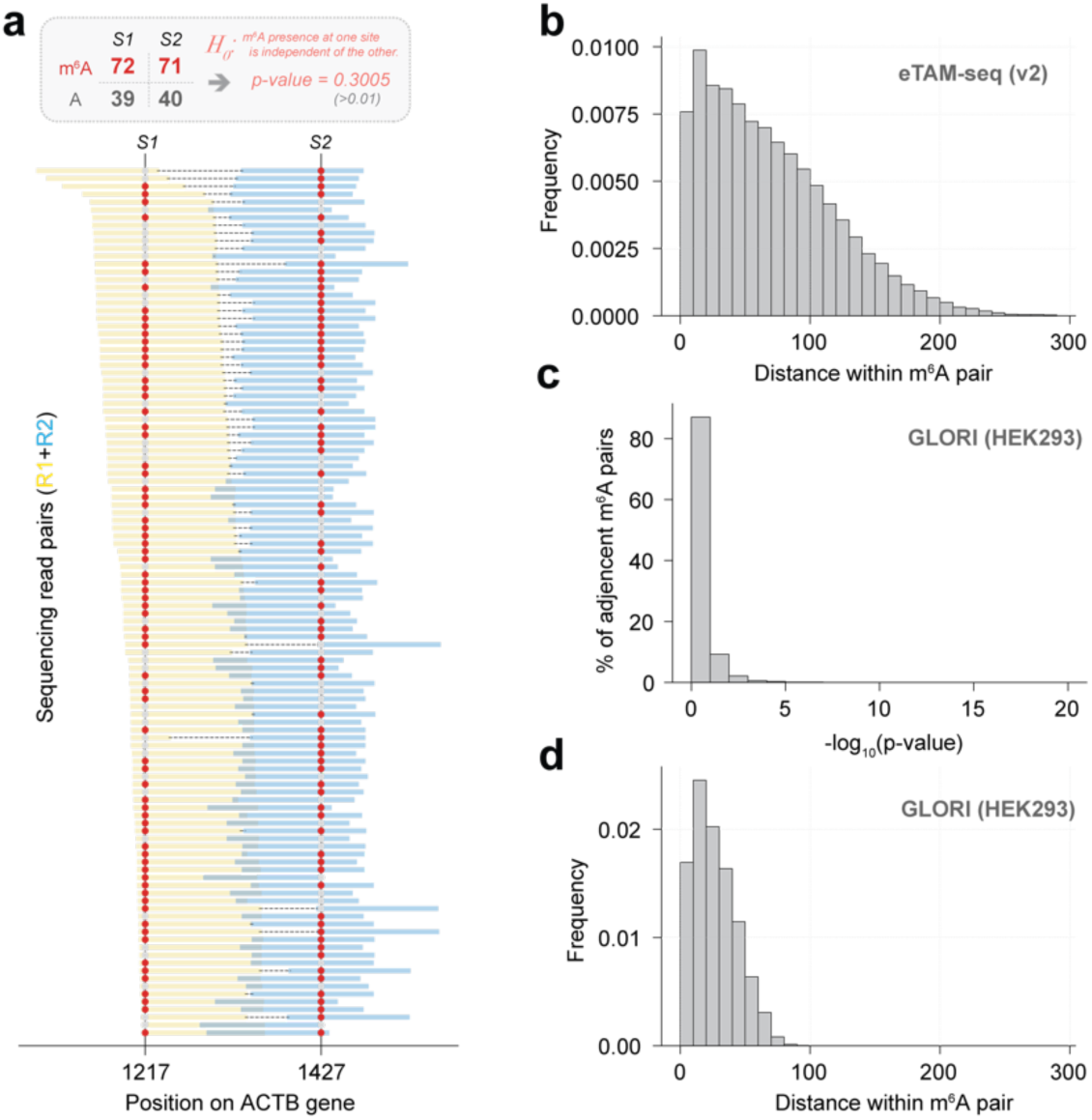
m^6^A co-occurrence in single transcript molecules. **a**. Example of an m^6^A pair on the *ACTB* gene, highlighting sites at positions 1217 and 1427. Sequencing reads simultaneously covering both sites are shown with paired-end reads connected by dashed lines. m^6^A signals are marked with red dots, and unmodified A signals with silver dots. **b**. Distance distribution of m^6^A pairs analyzed using eTAM-seq v2. **c**. Distribution of *P*-values from significance tests on co-occurrence for m^6^A pairs detected by GLORI. A total of 99,226 m^6^A sites (modification ratio >20%) formed 747,265 pairs; 29,373 pairs had at least 10 spanning reads, with 28,313 (96%) exhibiting *P*-values ≥0.01. **d**. Distance distribution of m^6^A pairs analyzed using GLORI. eTAM-seq-v2 HeLa replicate 1 was analyzed in **a**-**b**; GLORI HEK293T replicate 1 (ligation-based, SRR21356251) was analyzed in **c**-**d**.

**Extended Data Figure 5.**
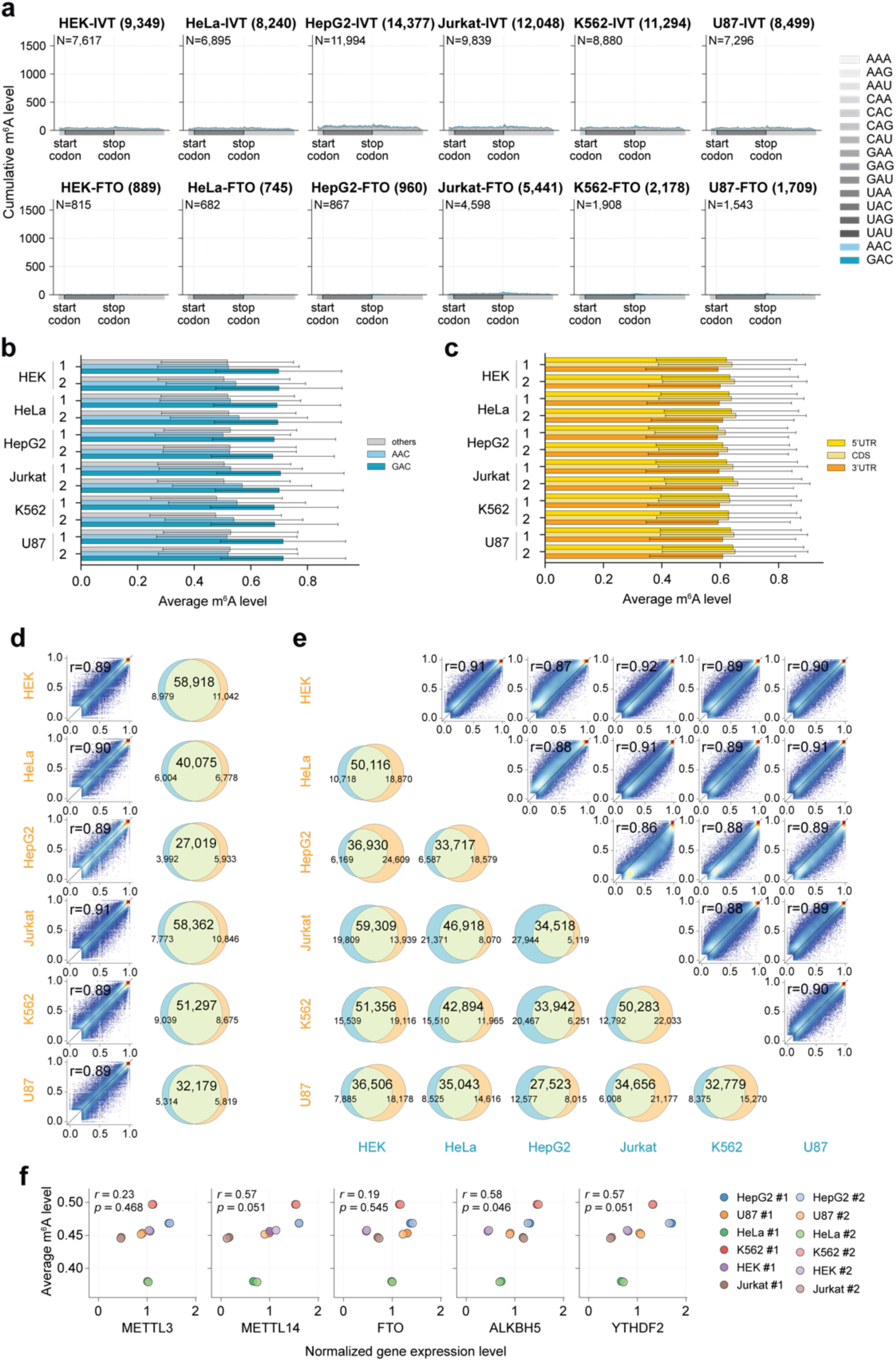
m^6^A profiling in six human cell lines. **a**. Metagene plots showing distribution of conversion-resistant A sites across transcript regions for control IVT and FTO-treated samples. **b**. Bar plots showing average m^6^A signals detected in GAC, AAC, and other NAN motifs. **c**. Bar plots showing average m^6^A levels detected across different transcript regions (5’ UTR, CDS, and 3’ UTR). **d**. Correlation of methylation levels between biological replicates for each of the six human cell lines. Pearson correlation coefficients (r) are indicated. Venn diagrams showing overlapping sites are provided on the right. **e**. Scatter plots showing pairwise correlation of methylation levels and Venn diagrams showing overlapping sites across six human cell lines. Methylation levels were averaged from two replicates to make the plots. Pearson correlation coefficients (r) are indicated. **f**. Scatter plots showing the correlation between global methylation levels and normalized gene expression levels of individual m^6^A writers (*METTL3*, *METTL14*), erasers (*FTO*, *ALKBH5*), and reader *YTHDF2* across six human cell lines.

**Extended Data Figure 6.**
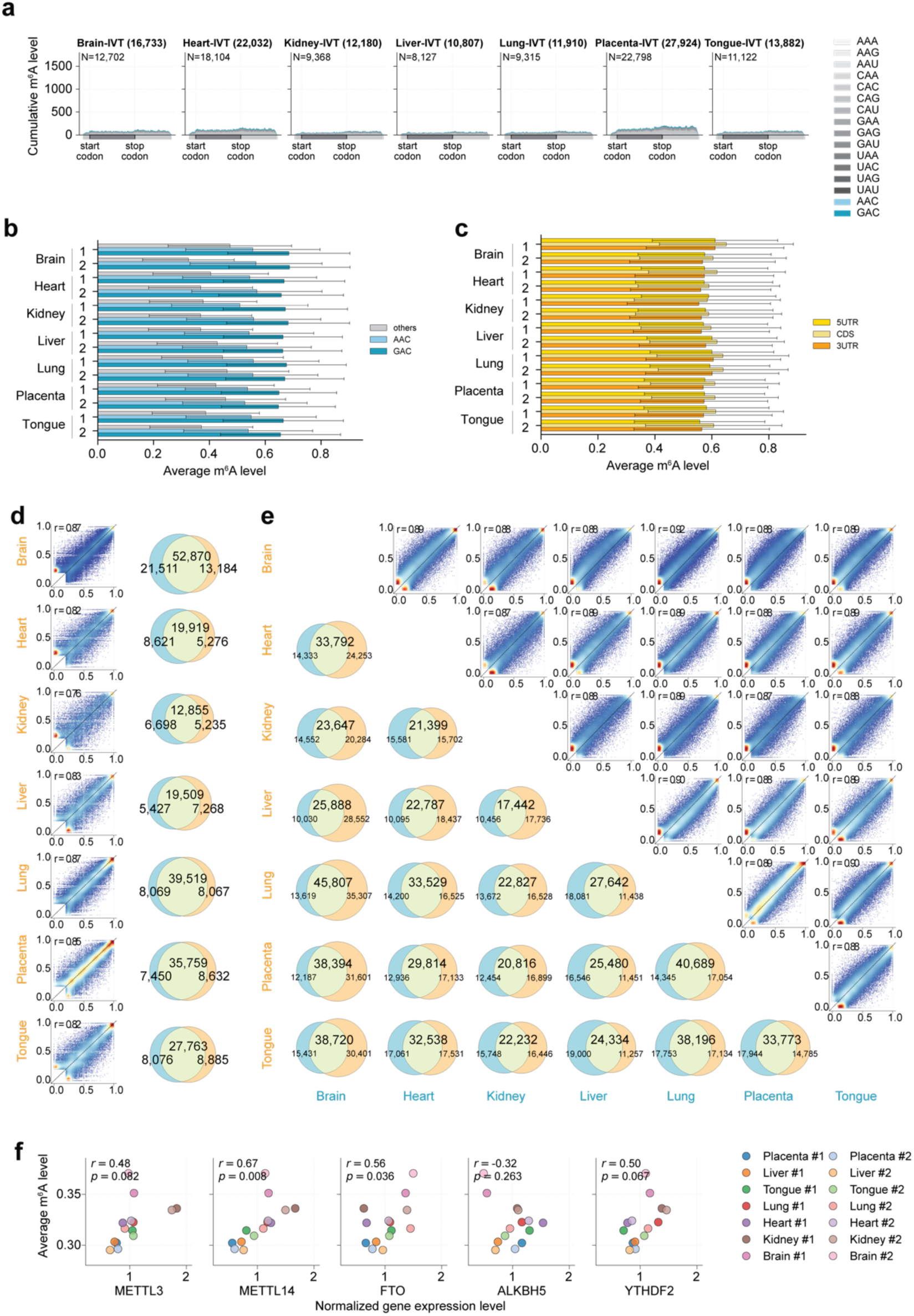
m^6^A profiling in seven embryonic mouse organs. **a.** Metagene plots showing distribution of conversion-resistant A sites across transcript regions for IVT controls. **b**. Bar plots showing average m^6^A signals detected in GAC, AAC, and other NAN motifs. **c**. Bar plots showing average m^6^A levels detected across different transcript regions (5’ UTR, CDS, and 3’ UTR). **d**. Correlation of methylation levels between biological replicates for each of the seven mouse tissues. Pearson correlation coefficients (r) are indicated. Venn diagrams showing overlapping sites are provided on the right. **e**. Scatter plots showing pairwise correlation of methylation levels and Venn diagrams showing overlapping sites across seven mouse tissues. Methylation levels were averaged from two replicates to make the plots. Pearson correlation coefficients (r) are indicated. **f**. Scatter plots showing the correlation between global methylation levels and normalized gene expression levels of individual m^6^A writers (*METTL3*, *METTL14*), erasers (*FTO*, *ALKBH5*), and reader *YTHDF2* across seven mouse tissues.

**Extended Data Figure 7.**
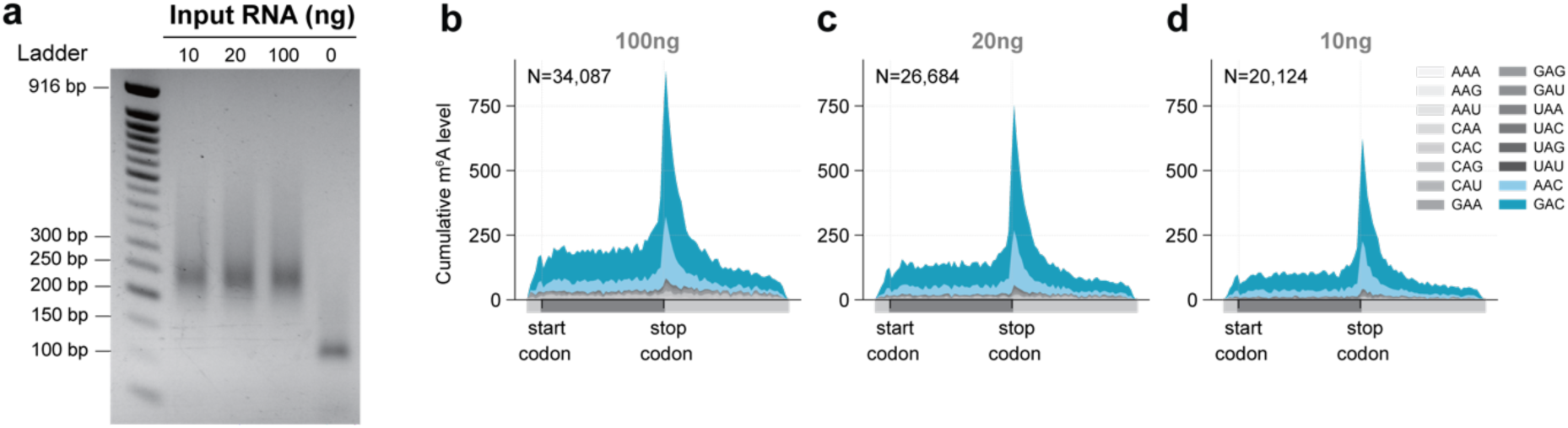
Performance of low-input eTAM-seq-v2. **a.** Agarose gel analysis of low-input eTAM-seq-v2 libraries constructed using 10/20/200 ng total RNA before size selection. **b**-**d.** Metagene plots showing the distribution of m^6^A sites across mRNA transcript regions detected using 100 (**b**), 20 (**c**), and 10 (**d**) ng of total RNA.

