## Supplementary Information for "Advanced eTAM-seq enables high-fidelity, low-input *N*^6^-methyladenosine profiling in human cells and embryonic mouse tissues"

**Supplementary Table**

**Supplementary Table 1** | Library quality assessment

**Supplementary Table 2** | $m^6A$ sites detected in human cell lines

**Supplementary Table 3** | $m^6A$ sites detected in mouse tissues

**Supplementary Table 4** | $m^6A$ sites detected in low-input samples

Supplementary Tables 2-4 are available for download at the Gene Expression Omnibus (GEO) under accession numbers GSE294681 (token: qfajakmcbjgnfux), GSE294682 (token: sbudamumzzmtpux) and GSE294683 (token: avajocsmjzylbkp).

**Supplementary Note 1** Modeling to distinguish $m^6A$ signals from sequence-dependent background

In negative detection methods, such as bisulfite sequencing for 5mC and eTAM-seq/GLORI for RNA $m^6A$ modifications, any insufficient conversion is erroneously interpreted as a modification, leading to false positives. Thus, conversion efficiency is critical.

Several factors influence conversion efficiency:
**Motif sequence context**: Deamination reaction efficiency is sequence-dependent, and different motifs exhibit varying conversion rates.
**Local sequence structure**: Structures such as RNA G-quadruplexes (rG4s) can significantly hinder enzyme recognition and lower conversion efficiency. Guanine-cytosine content (**GC%**, noted hereafter) is a major factor affecting local structure formation.
**RNA 3D structure**: Long-range stem-loop regions can impede conversion. Although this issue is largely mitigated by denaturation steps and the use of TadA enzyme at elevated reaction temperatures, further improvement can be achieved computationally by filtering out reads with constitutively unconverted sites. This approach leverages the knowledge that only approximately 0.5% of adenosine sites in human and mouse transcripts are modified as $m^6A$.

To address these challenges and enhance the specificity of $m^6A$ detection, we implemented a computational strategy involving initial read filtering followed by a sequence-context-aware modeling approach to distinguish true $m^6A$ signals from GC-dependent background noise.

### Step1: **Initial read processing and site-level data generation**

Before modeling background rates, raw sequencing reads were processed to generate site-level conversion information across the transcriptome:

- **Filtering reads with consecutive unconverted sites**: Reads exhibiting stretches of potentially incomplete conversion, defined as having 3 or more consecutive unconverted adenosine (A) sites, were removed. This step helps eliminate reads likely affected by broader technical issues rather than specific local modification.
- **Read trimming**: To mitigate potential biases from adapters or non-templated nucleotides sometimes added during RT, the first and last 2 nucleotides of each read were trimmed.
- **Base counting**: After filtering and trimming, reads were mapped to the reference transcriptome. For every adenosine (A) site covered reads, the number of reads showing conversion (G) at that site and the number showing non-conversion (remaining A) were counted across all mapped reads. This generated the raw input data: per-site unconverted counts (**u**) and total coverage (**d**).

### Step2: **Modeling GC-content dependent unconverted ratios**

The core of our computational refinement involves modeling the observed unconverted ratio (**r=u/d**) as a function of local GC content, specific to each sequence motif context.

**2.1. Flanking GC content calculation:** For each A site, the local GC content was calculated within a 20-nucleotide window flanking the site: 10-nt upstream and 10-nt downstream. The number of 'G' or 'C' bases within this 20-nt window was counted and divided by 20 to obtain the GC fraction, denoted as **x**.

**2.2. Data aggregation for modeling**
Site-level data was aggregated for each experimental library:

- **Calculating ratios:** Within each library, the site-level unconverted ratio (**r=u/d**) was calculated.
- **GC binning:** To reduce noise and handle sparse data at extreme GC values, sites with $x<0.2$ were consolidated into a single 'low GC' bin, and sites with $x>0.8$ were consolidated into a single 'high GC' bin.

- **Grouping and average**: Sites were further grouped based on their GC bin for each 3-nt sequence context (NAN, where N is any base, centered on the target 'A'). Within each resulting NAN+GC group, the average ratio (denoted as $\bar{\mathbf{r}}$) and average GC (denoted as $\bar{\mathbf{x}}$) were calculated. This aggregation yields a set of representatives ($\bar{\mathbf{x}}$, $\bar{\mathbf{r}}$) data points for each motif, suitable for curve fitting.

### 2.3. Mathematical framework

Analysis revealed that the background rate of apparent resistance to A-to-G increases exponentially with higher local GC content (**Extended Data Fig. 3a**). While the majority of adenosine sites reside in regions of moderate GC content (**Extended Data Fig. 3b**), this GC-dependent background effect, observed consistently across all eTAM-seq libraries (**Supplementary Note Figure 1**), could potentially bias $m^6A$ quantification.

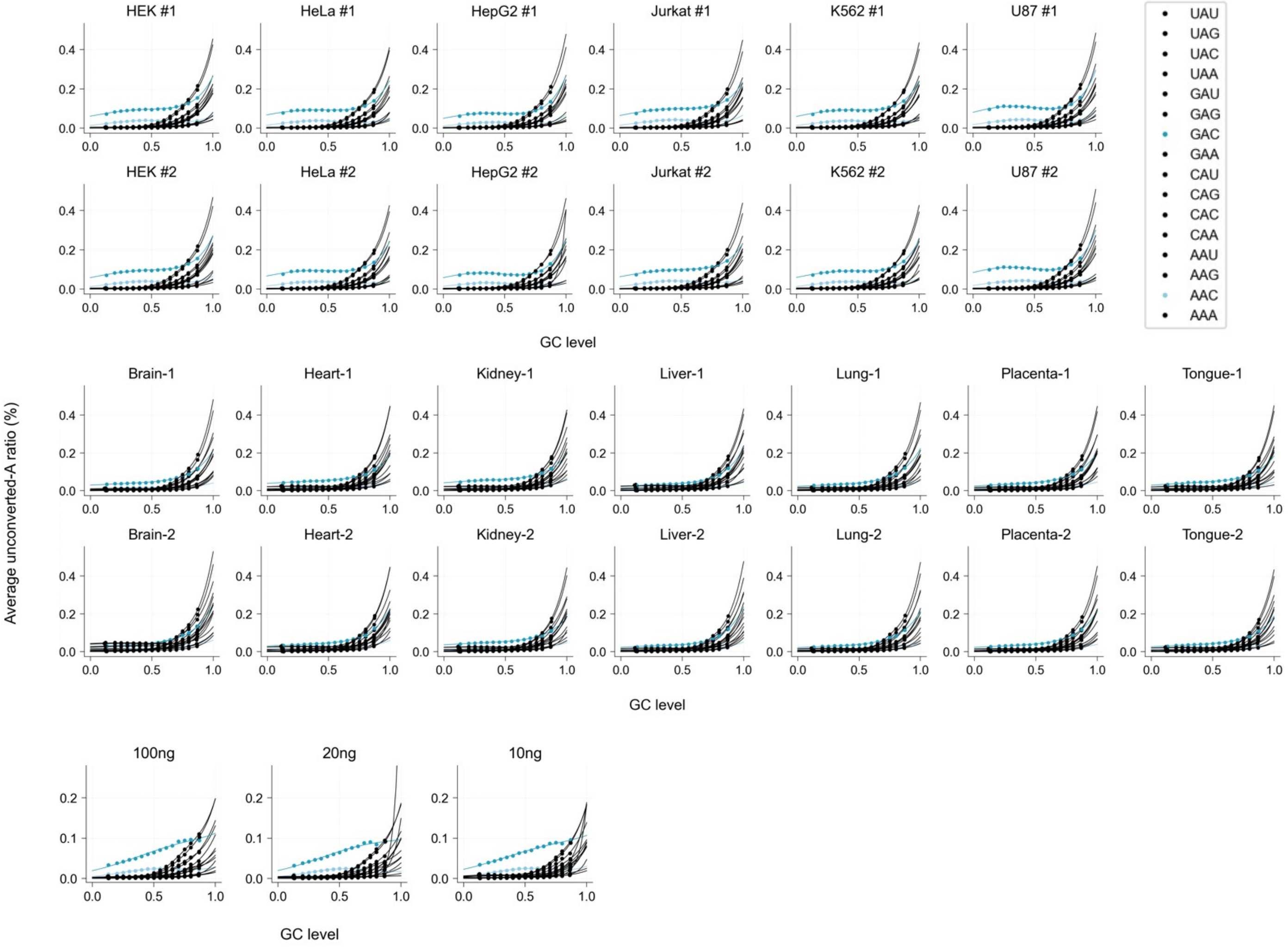


**Supplementary Note Figure 1**: Relationship between adenosine conversion resistance and local GC content.

To account for this bias, we developed a mathematical model describing the relationship between the aggregated average observed unconverted ratio ($\bar{\mathbf{r}}$) and the aggregated average local GC content ($\bar{\mathbf{x}}$) for specific sequence motifs. Initial analysis showed that a simple exponential function could approximate the trend for some motifs (e.g., CAC), but failed for known $m^6A$-containing motifs (e.g., AAC). These $m^6A$ motifs exhibited a distinct baseline unconverted ratio at low GC content, attributable to true $m^6A$ modification, and also showed a pattern where the apparent $m^6A$ signal varied with GC content (**Supplementary Note Figure 1**). This suggested that both the background conversion resistance and the $m^6A$ signal itself are functions of local GC content.

Therefore, the model incorporates two components:

- **Background rate:** Modeled as an exponential function $f(\overline{\mathbf{x}}, \mathrm{m}, \mathrm{n}) = m \cdot e^{m\overline{\mathbf{x}}}$, whereas m and n are two constant values for each motif.
- **m$^6$A signal component:** Modeled as a Gaussian-like function $g(\overline{\mathbf{x}}, \mathrm{o}, \mathrm{p}, \mathrm{q}) = o \cdot e^{-p(\overline{\boldsymbol{x}}-\boldsymbol{q})^2}$, wherease o, p, and q are three constant values for each motif.
- **Combined model:** The overall observed ratio y is modeled by integrating the background rate and the m$^6$A signal component: $h(\overline{\mathbf{x}}, \mathrm{m}, \mathrm{n}, \mathrm{o}, \mathrm{p}, \mathrm{q}) = g(\overline{\mathbf{x}}, o, p, q) + \left(1 - g(\overline{\mathbf{x}}, o, p, q)\right) \cdot f(\overline{\mathbf{x}}, \mathrm{m}, \mathrm{n})$
- **Parameter fitting procedure:** The five parameters $(\mathrm{m}, \mathrm{n}, \mathrm{o}, \mathrm{p}, \mathrm{q})$ of the combined model were determined for each motif characterizing its specific GC-dependent unconverted ratio profile.

### Step3: **Statistical validation and filter sites**

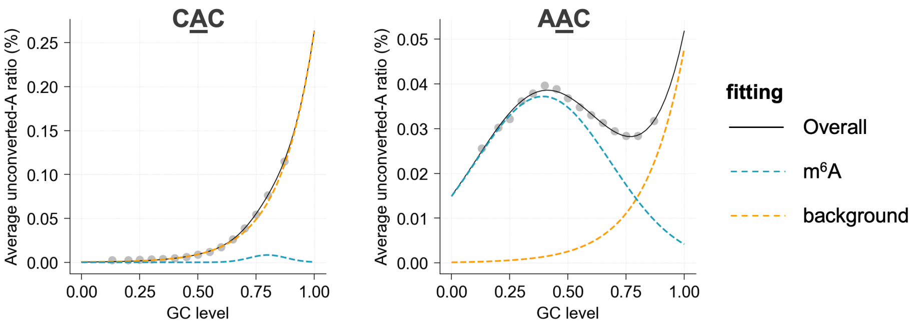


**Supplementary Note Figure 2**: Fitting of CAC and AAC motif from HeLa replicate 1 as example.

After model fitting (**Supplementary Note Figure 2**), the functions for GC-dependent background rate fitting, $f(\overline{\mathbf{x}}, \mathrm{m}, \mathrm{n})$ were extracted for each motif. Subsequently, each potential modification site was statistically evaluated and filtered to determine if its observed unconverted ratio significantly exceeded the predicted background level.

- **Background rate assignment:** First, the expected background rate (***b***) was calculated for every A site in the dataset. This was achieved by evaluating the fitted background function with fitted parameters $(\mathrm{m}, \mathrm{n})$ corresponding to the site's 3-nt motif for the specific library being analyzed, and a minimum background rate of 0.0001 was enforced for sites of too low background noise or ensure sufficient data in fitting.
- **Statistical Testing:** One-sample Chi-squared test was performed. This test compared the observed distribution of unconverted vs. converted reads at the site against the expected distribution under the null hypothesis that the site's true unconverted rate (***r***) is equal to the predicted background rate ***b***. Specifically, it tests if the observed distribution $[r, 1-r]$ is significantly different from the expected distribution $[b, 1-b]$. Low p-values thus indicate that the observed ratio is significantly higher than the modeled background.
- **Site Filtering:** For each site within each library, several filtering steps were applied to call m$^6$A sites:
    - sequencing depth (d) >10
    - observed unconverted ratio (r) ≥0.2
    - significance threshold (p-value) <0.0001
